# Candelabrum cells are molecularly distinct, ubiquitous interneurons of the cerebellar cortex with specialized circuit properties

**DOI:** 10.1101/2021.04.09.439172

**Authors:** Tomas Osorno, Stephanie Rudolph, Tri Nguyen, Velina Kozareva, Naeem Nadaf, Evan Z. Macosko, Wei-Chung Allen Lee, Wade G. Regehr

**Author notes:** Equal contribution (Tomas Osorno, Stephanie Rudolph). Albert Einstein College of Medicine, New York, NY 10461. Correponding author.

## Abstract

To understand how the cerebellar cortex transforms mossy fiber (MF) inputs into Purkinje cell (PC) outputs, it is vital to delineate the elements of this circuit. Candelabrum cells (CCs) are enigmatic interneurons of the cerebellar cortex that have been identified based on their morphology, but their electrophysiological properties, synaptic connections, and function remain unknown. Here we clarify these properties using electrophysiology, snRNA sequencing, *in situ* hybridization, and serial electron microscopy. We find that CCs are the most abundant PC layer interneuron. They are GABAergic, molecularly distinct, and present in all cerebellar lobules. Their high resistance renders CC firing highly sensitive to synaptic inputs. CCs are excited by MFs and granule cells, and strongly inhibited by PCs. CCs in turn inhibit molecular layer interneurons, which leads to PC disinhibition. Thus, inputs, outputs and local signals all converge onto CCs to allow them to assume a unique role in controlling cerebellar output.

## Introduction

The cerebellum controls fine motor coordination, motor learning, eye movements, balance, and cognitive-affective functions, among many other behaviors that depend on prediction (Schmahmann, 2019; Sokolov et al., 2017; Strick et al., 2009; Wang et al., 2014). The cerebellar cortex is thought to contribute to these behaviors by transforming multimodal sensory and higher-order inputs from mossy fibers (MFs) into Purkinje cell (PC) outputs. Within the cerebellar cortex, few MFs excite many granule cells (GCs) that in turn form weak synapses onto a moderate number of PCs. Inhibitory interneurons contribute to information processing in all stages. Golgi cells (GoCs) regulate GC excitability and MF integration in the input layer (D’Angelo et al., 2013; Duguid et al., 2015; Mitchell and Silver, 2000). Molecular layer interneurons (MLIs) inhibit PCs to control the output of the cerebellar cortex (Jorntell et al., 2010; Kim and Augustine, 2020). Although the simplicity of this circuit holds great promise for determining how circuit level computations contribute to behavior, the cerebellum appears to contain additional elements that are neglected in current circuit models. It is particularly important to delineate these cell types for the cerebellar cortex, where just seven types of interneurons have been identified, compared to approximately 50-60 types of inhibitory interneurons in cortical areas such as V1, ALM and M1 (Adkins et al., 2020; Bakken et al., 2020; Tasic et al., 2018; Yao et al., 2020). It is therefore mandatory to identify and characterize elusive circuit elements to better understand how they contribute to computations within the cerebellar circuit and ultimately to behavior.

The candelabrum cell (CC) is the most mysterious neuron of the cerebellar cortex. CCs were identified in 1994 based on their distinctive light-level morphology (Laine and Axelrad, 1994). CCs have a small cell body near the Purkinje cell layer (PCL), dendrites that extend to the surface of the molecular layer, and beaded axons that make numerous local synapses within the molecular layer. Nothing else was known about CCs because there had been no electrophysiological recordings, and no molecular markers had been identified. In our recent snRNAseq study we identified three types of molecularly distinct inhibitory interneurons in the cerebellar cortex that are neither MLIs or GoCs (Kozareva et al., 2020). These findings have the potential to provide insight into the molecular properties of CCs. However, although it seems likely that one of these cell types corresponds to CCs, the link between these molecularly defined populations and CCs is not known.

Here, we use a combination of mouse genetics, single nucleus sequencing data, *in situ* hybridization, electrophysiology and serial EM reconstructions to characterize CCs. We identified a transgenic mouse line in which CCs are fluorescently labeled that allowed us to record from CCs, and we found that they are highly sensitive to small depolarizations. We found that CCs are a molecularly distinct population of GABAergic interneurons present in all cerebellar lobules, and they are almost as prevalent as GoCs. Electrophysiology and serial electron microscopy (EM) reconstructions show that PCs inhibit CCs. CCs in turn inhibit MLIs, leading to the disinhibition of PCs. Therefore, CCs are part of an inhibitory ring between MLIs, PCs and CCs. Their ability to weigh MF inputs, PC outputs and GC activity, indicates that the function of CCs is distinct from that of MLIs and GoCs. CCs must therefore be considered an important interneuron of the cerebellar cortex that need to be incorporated into circuit models to gain a full understanding of cerebellar processing.

## RESULTS

### Morphological and electrophysiological properties of Candelabrum cells

Recording from CCs is vital to clarifying their role in the cerebellar circuit. This had not been done previously because CCs are located in the PCL and cannot be readily distinguished from the more abundant granule cells, MLIs and Bergmann glia that have approximately the same cell body size and position. In our experience, it was also difficult to identify CCs using mice that express GFP in GABAergic glutamate decarboxylase 67-positive (GAD67-GFP) neurons because CCs are typically obscured by much brighter PCs. Fortunately, we found that the oxytocin receptor Cre (Oxtr-Cre) line crossed to a *flox* tdTomato reporter line (Ai14) provided a means of labelling CCs. In Oxtr-Cre x Ai14 mice, small interneurons of the PCL (PLIs) were labelled that had processes extending into the molecular layer (**Figure 1A)**. MFs and a small number of MLIs and PCs were also labelled, but they were readily distinguished from PLIs based on their position, shape and size.

**Figure 1.**
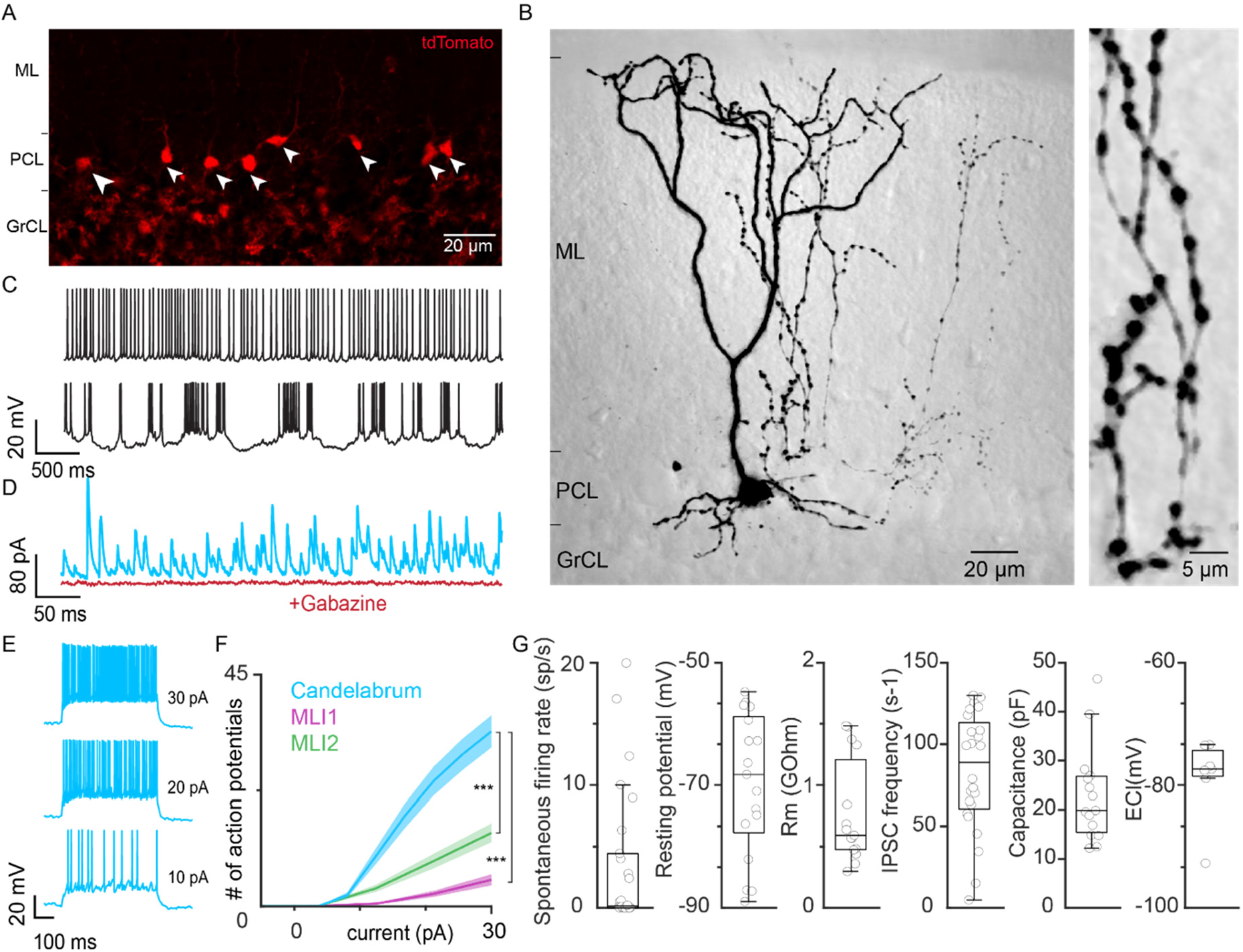
Determining the morphological and electrophysiological properties of candelabrum cells using Oxtr-Cre x Ai14 (tdTomato) mice. A. Cerebellar cortex of Oxtr-Cre x Ai14 showing small cells in the PCL (white arrows) and MFs labeled with tdTomato (red) in the granule cell layer. B. 2-photon image of a labeled candelabrum cell. Image to the right is an expanded image of the axon. C. Spontaneous activity in a regular firing (top) and burst firing (bottom) candelabrum cell. D. Spontaneous IPSCs recorded in a candelabrum cell in the absence of synaptic blockers (light blue) and after wash-in of the GABA_A_R antagonist gabazine (red). E. Examples of firing evoked by current injection of 10, 20 and 30 pA. F. Average input-output curve of candelabrum cells, compared to the input-output curve of two types of molecular layer interneurons, MLI1 and MLI2 (data from Kozareva et al. 2020). p=2.1e-18 (for slope MLI1/CC), p= 7.5e-10 (for slope MLI2/CC), generalized mixed effects model. (MLI1: n=22; MLI2: n=20; CC: n=18). G. Summary of basic electrophysiological properties of candelabrum cells. (n=26,17,15, 24,15, 7 cells)

We determined the morphology of tdTomato+ PLIs by imaging cells that were filled with Alexa-594 during whole-cell recordings. All labelled cells exhibited the characteristic morphology of CCs, with small cell bodies (~10 μm in diameter) located in the PCL, one or two primary dendrites that ascend to the pia and branch within the upper molecular layer, several secondary dendrites that extend to the upper part of the granular layer, and a beaded axon emanating from the proximal primary dendrite that branches and extends in the molecular layer (**Figure 1B**). The stereotyped morphology of these neurons was in concordance with the original description of CCs (**Figure 1S1**)(Laine and Axelrad, 1994), and we conclude that we are able to consistently record from CCs in Oxtr-Cre x Ai14 mice.

In cell-attached recordings and whole-cell current clamp recordings, CCs were mostly silent, although some fired spontaneously, with varying frequencies and regularity (**Figure 1C, G**). In voltage-clamp recordings, high-frequency spontaneous IPSCs were observed that were blocked by the GABA_A_ receptor antagonist gabazine (10 μM; **Figure 1D, G**). Input-output curves show that CCs are extremely sensitive to current injection, with current steps as small as 5 pA evoking spikes (**Figure 1E, F**). This contrasts with the two types of MLIs, MLI1 and MLI2, that are much less sensitive to current injection (**Figure 1F**) (Kozareva et al., 2020). A summary of the basic electrophysiological properties of CCs (**Figure 1G**) suggests that the most distinctive characteristics of CCs are 1) high frequencies of sIPSCs that are absent in GoCs and MLIs, and 2) high excitability that far exceeds the excitability of MLIs.

### Molecular insights into interneurons of the cerebellar cortex

Our previous molecular characterization of cerebellar cell types has the potential to provide insight into the molecular identity of CCs. We used single nucleus RNA sequencing (snRNAseq) data to molecularly characterize neurons of the adult cerebellar cortex and identified eight clusters of inhibitory interneurons (Kozareva et al., 2020). In addition to molecular layer interneurons (MLIs that can be subdivided into MLI1_1, MLI1_2, and MLI2), and GoCs (Golgi1 and Golgi2), we identified three types of inhibitory interneurons that were neither GoCs or MLIs, that we referred to as Purkinje layer interneurons (PLI) PLI1, PLI2, and PLI3 (**Figure 2A**). However, the correspondence between these molecularly defined cell types and known cell types of cerebellar interneurons is unclear.

**Figure 2.**
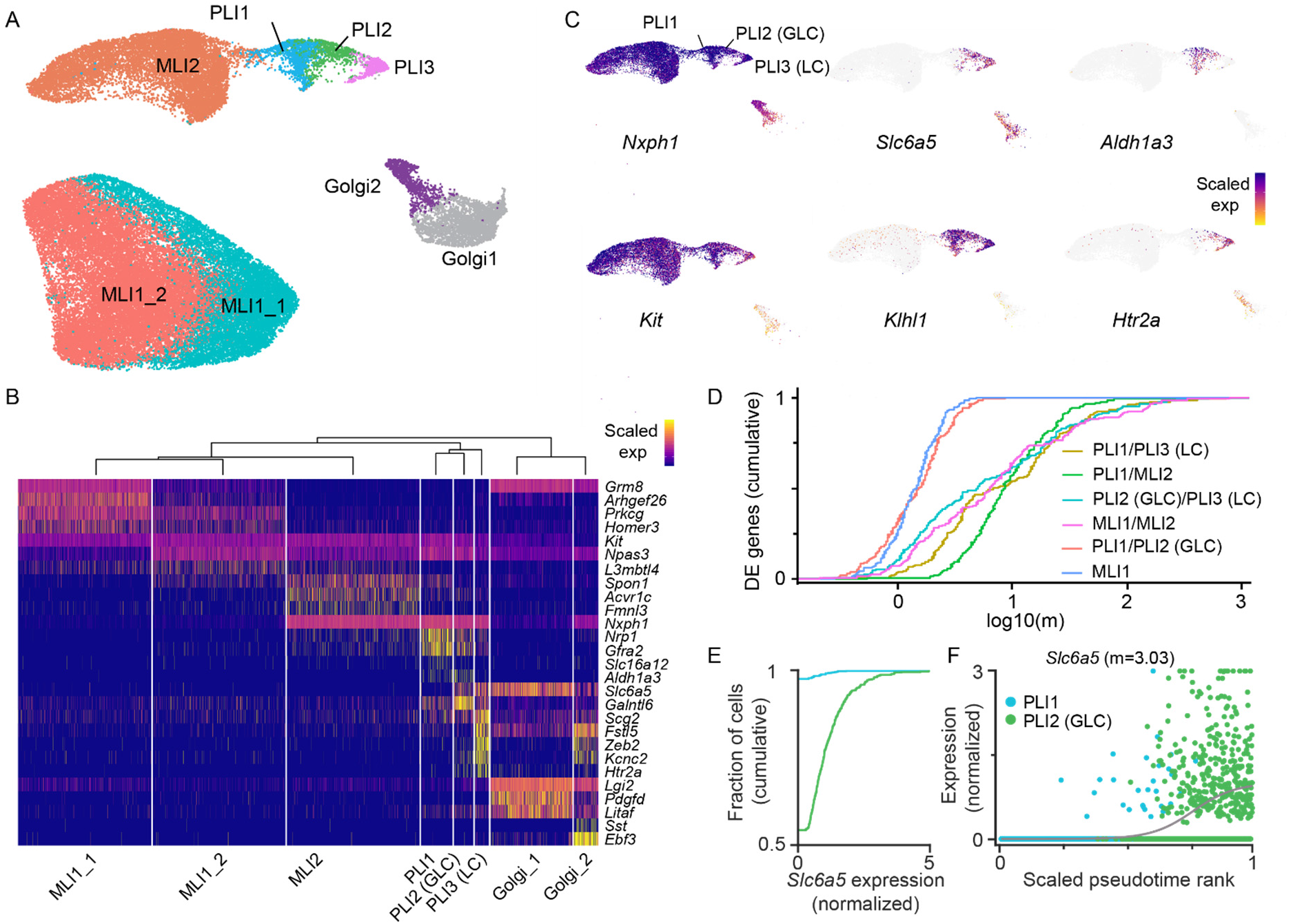
Characterization of cerebellar inhibitory interneurons with snRNA seq. A. UMAP plot of cerebellar inhibitory interneurons. MLI1 and ML2 are types of molecular layer interneurons, and MLI1 can be further subdivided into MLI1_1 and MLI1_2. Golgi1 and Golgi2 are types of Golgi cells. Purkinje cell layer interneurons are divided into PLI1, PLI2 and PLI3. B. Scaled RNA expression of key genes in cerebellar inhibitory interneurons C. UMAP plots of *Nxph1, Slc6a5, Aldh1a3, Kit, Klhl1* and *Htr2a* expression in MLI2, PLIs and Golgi2 cells. Colormap maximum values were adjusted for each gene (*Nxph1:* 5.5, *Slc6a5/Htr2a:* 2.5, *Aldh1a3:* 2, *Kit/Klh1:* 3.5) D. Cumulative distributions of *m* values from logistic curve fits (as in F) for top differentially expressed (DE) genes among aggregated combinations of the indicated cell types. E. Cumulative distributions of *Slc6a5* expression in PLI1 and PLI2 (GLC). F. Expression of *Slc6a5* as a function of pseudotime in PLI1 and PLI2 (GLC). The grey curve is a logistic fit, with the maximum slope value of the fit (m) indicated. 14 cells with normalized expression greater than 3 are plotted at 3.

A summary of the expression of key genes showed that, as is often the case, it is not always possible to delineate a cell type with a single gene. Rather, a combination of genes is required to demarcate distinct interneuron types, including PLI1, PLI2 and PLI3 (**Figure 2B,C**). The expression of *Htr2a* that encodes the 5-HT2A receptor, and *Slc6a5* that encodes the glycine transporter, by PLI3 suggests that this cell type corresponds to Lugaro cells (LCs) that are glycinergic and excited by serotonin (Dieudonne and Dumoulin, 2000). LCs are GABAergic/glycinegic inhibitory PLIs with a distinctive fusiform soma that are inhibited by PCs, and that locally inhibit GoCs and MLIs, and send long-range axons to unknown targets (Dieudonne, 2001; Dieudonne and Dumoulin, 2000; Geurts et al., 2002; Geurts et al., 2001; Laine and Axelrad, 1996; Palay and Chan-Palay, 1974; Sahin and Hockfield, 1990; Simat et al., 2007). PLI2s also express *Slc6a5*, which suggests that they correspond to globular cells (GLCs) that are glycinegic cells located near or below the PCL and are inhibited by PCs (Hirono et al., 2012; Laine and Axelrad, 2002). This raised the possibility that CCs might correspond to the remaining cell type, PLI1. PLI1s can be uniquely identified by the combined expression of *Nxph1, Aldh1a3*, and the lack of *Slc6a5* expression. *Nxph1* encodes the secreted protein Neurexophilin-1, *Slc6a5* encodes the glycine transporter GlyT2, and *Aldh1a3* encodes an aldehyde dehydrogenase that oxidizes all-trans-retinal to retinoic acid. Expression of *Oxtr* was not detected in the PLI1 population (data not shown).

We next applied a method we developed previously (Kozareva et al., 2020) to characterize the relative discreteness versus continuity across the clustered populations. Briefly, we fit logistic curves to the expression of differentially expressed genes along the pseudotime expression trajectory constructed across different pairs of interneuron populations. We then determined the maximum slope value of the fit, *m*, for 150 highly variable genes (**Figure 2D**). High values of *m* indicate sharp transitions and suggest discrete cell types, whereas low values indicate smooth transitions and gradations in cell class. PLI1s were distinct from LCs (PLI3s) and MLI2 cells, and LCs were distinct from GLCs (PLI2s). These aggregate distributions were similar to those of the aggregate distributions of the discrete cell types MLI1 and MLI2. PLI1s and GLCs were more similar to each other and exhibited an aggregate distribution similar to MLI1, which consists of MLI1_1 and MLI1_2 subtypes. These findings indicate that PLI1s are more similar to GLCs than any other type of neuron.

We assessed whether it would be possible to overcome the molecular similarity of PLI1s and GLCs in order to discriminate between these cell types. *Slc6a5* was detected in 48% (341/735) of GLCs, but only 2% of PLI1s (28/1177 cells) (**Figure 2E**). We also plotted *Slc6a5* expression as a function of pseudotime and found that this was approximated with a logistic fit with a maximum slope, *m*, of 3.03 (**Figure 2F**). Thus, *Slc6a5* expression provides promising means for discriminating between PLI1s and GLCs.

### Candelabrum cells are prevalent and widely distributed in the cerebellar cortex

We used single molecule FISH (smFISH) in sagittal sections of adult cerebellum vermis to provide insight into PLI1 and determine whether they corresponded to CCs (**Figure 3**). We used three probes to analyze the distribution of different types of interneurons in wildtype animals (**Figure 3A-E**), and used two of these probes to characterize tdTomato+ neurons in Oxtr-Cre x Ai14 mice (**Figure 3F-J**). These experiments established that PLI1 neurons correspond to the tdTomato+ neurons in Oxtr-Cre x Ai14 mice that correspond to CCs. For clarity, we therefore refer to PLI1 as CCs in **Figure 3**.

**Figure 3.**
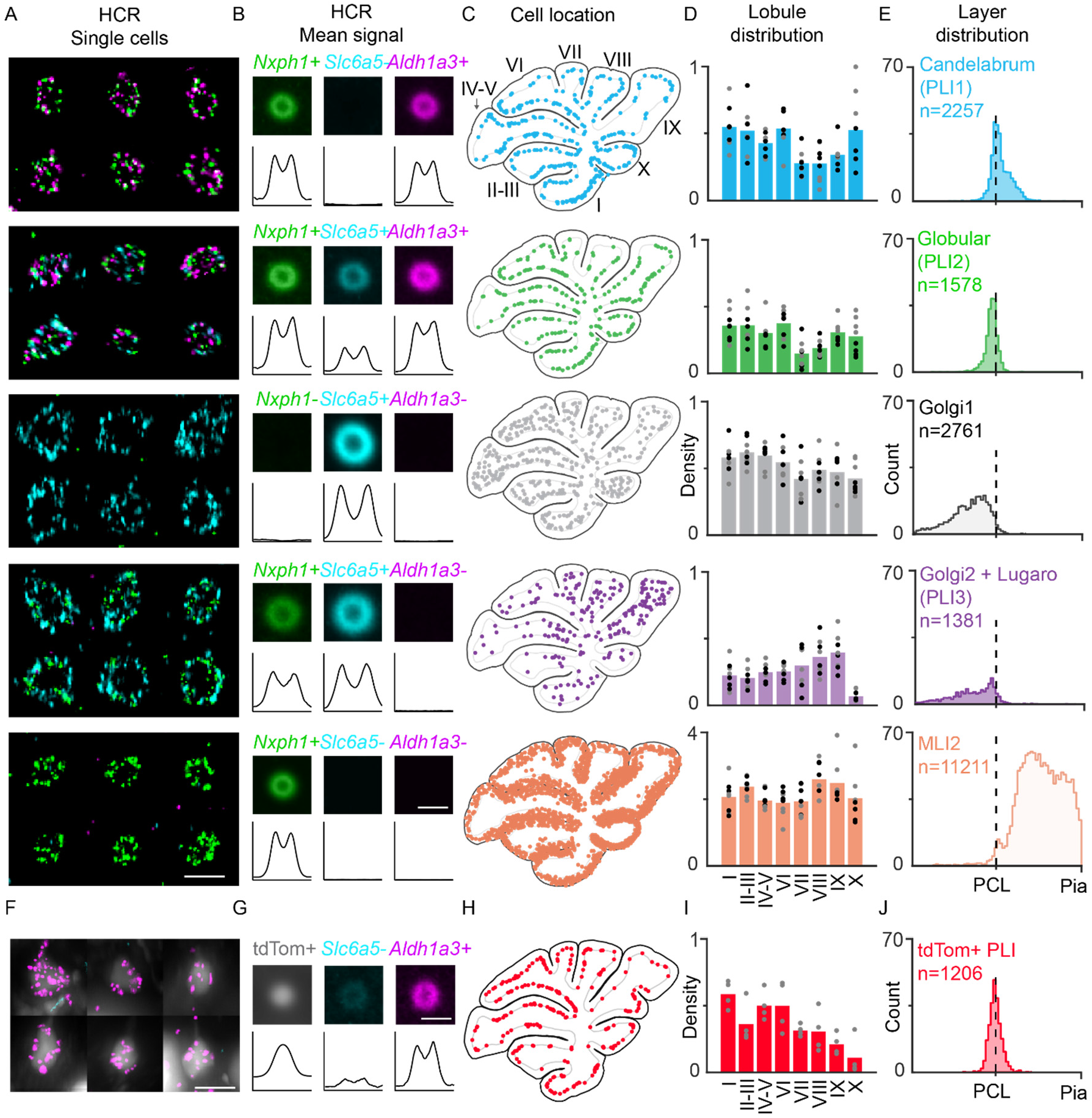
Identification of different types of interneurons in the cerebellar cortex with fluorescent in situ hybridization. A. Example labeling of single cells for five different classes of cells with probes for *Nxph1, Aldh1a3* and *Slc6a5* RNA (top to bottom):

Candelabrum cell (PLI1): *Nxph1+, Slc6a5-, Aldh1a3+*
Globular cell (PLI2): *Nxph1+, Slc6a5+, Aldh1a3+*
Golgi1: *Nxph1-, Slc6a5+, Aldh1a3-*
Golgi2 + Lugaro cells (PLI3): *Nxph1+, Slc6a5+, Aldh1a3-*
MLI2: *Nxph1+, Slc6a5-, Aldh1a3-* B. Mean fluorescence signals for three probes for the five classes of cells. Lower plots show intensity profiles for each channel. C. Locations of cells in a single slice for five classes of cells. D. Average densities of different types of cells in different lobules. Each point is from a different slice, male (black) female (gray). E. Histograms of the locations of each cell type relative to the center of the PCL, and the top of the molecular layer (Pia). n=8 slices. F-J Same as A-E for tdTom+ PLIs in Oxtr-Cre x Ai14 mice. Four slices are summarized in I and J.

We used probes for *Nxph1, Slc6a5* and *Aldh1a3* (**Figure 2C**, *top*), in order to determine the position and quantify the fluorescence for each labelled cell in cerebellar slices (**Figure 3S1**). Fluorescence images of different types of cells (**Figure 3A**), and average images and line profiles (**Figure 3B**) are shown for the 3 labels for each cell type. The combined expression of these 3 markers allowed us to identify five populations of cells: CCs (*Nxph1+, Aldh1a+, Slc6a5-*), GLCs (*Nxph1+, Aldh1a3+, Slc6a5+*), Golgi1 cells (*Nxph1-, Aldh1a3-, Slc6a5+*), a mixed population that contains Golgi2 cells and LCs (*Nxph1+, Aldh1a3-, Slc6a5+*), and MLI2s (*Nxph1+, Aldh1a3-, Slc6a5-*). We also determined the spatial distributions of the different cell types (**Figure 3C**), quantified their numbers in different lobules (**Figure 3D**), and determined their locations relative to the PCL (**Figure 3E**). Cell distributions were determined in a total of eight slices from a male and a female mouse, and there were no obvious sex-dependent differences in the distribution of cells (**Figure 3S2**). CCs were located in all lobules of the vermis, although their density was lower in lobules VII, VIII, IX (**Figure 3C, 3D, 3S2**, *light blue*). CCs were clustered around the PCL, with a fraction of CCs extending into the lower third of the molecular layer (**Figure 3E**, *light blue*). CCs were remarkably abundant, with the ratio of the number of given cell type and the number of CCs being: 0.70 GLCs, 1.22 Golgi1, 0.61 for combined Golgi2 and LCs, and 4.96 for MLI2. These ratios are consistent with estimates based on snRNAseq data, where the ratios were 0.62 (735/1177) for GLCs and 0.45 (531/1177) for LCs (**Figure 3 S3**). Thus, there are approximately the same number of PLI interneurons as GoCs, and CCs are the most abundant PLI interneuron.

Each type of interneuron displayed a characteristic distribution. The distribution of GLCs was most similar to that of CCs (**Figure 3C, 3D, 3S2**, *green*). GLCs were clustered around the PCL, although unlike CCs, GLCs were not present in the molecular layer but extended into the upper part of the granular layer (**Figure 3E**, *green*). Golgi1 cells were found in all lobules, and they were restricted to the granule cell layer (**Figure 3C, 3D, 3S2**, *grey*). Although our labelling strategy was focused on identifying CCs, rather than discriminating between LCs and Golgi2 cells, the combined labelling of these two cell types is informative (**Figure 3C, 3D, 3S2**, *light purple*). The density of cells shows a gradient in the different lobules, peaks in lobule IX, and is extremely low in lobule X. The small peak in cells near the PCL likely reflects LCs, which are expected to be present at much lower densities than other PLIs. As expected, MLI2s are present at higher densities than the other cell types and are restricted to the molecular layer (**Figure 3C, 3D, 3S2**, *orange*).

In Oxtr-Cre x Ai14 mice, many small neurons of the PCL were labelled with tdTomato along with a small number of MLIs and PCs (**Figure 3S4**). The labeled MLIs and PCs were readily distinguished from small PLIs based on their position, shape and size. We found that tdTomato+ PLIs expressed *Aldh1a3*, but *Slc6a5* expression was low or absent (**Figure 3F, G. Figure 3S4**). The distribution of tdTomato-labelled PCL neurons (**Figure 3C-E**) was similar to that of CCs (PLI1) with the exception that there were fewer cells observed in lobule X (**Figure 3I**), and fewer cells in the lower molecular layer (**Figure 3J**). Thus, the combination of the tdTomato+ labelling of PLIs in Oxtr-Cre x Ai14 mice, our fluorescent labelling of these cells (**Figure 1**), and *in situ* labelling with probes identified from our single cell characterization, establishes that PLI1 correspond to CCs.

### Excitatory synapses onto Candelabrum cells

Neurons that extend dendrites into the molecular layer, including MLIs, GoCs and PCs, are all excited by the axons of granule cells, the parallel fibers (PF). We therefore tested whether granule cells also excite CCs. We used an extracellular electrode to excite parallel fibers with pairs of stimuli and recorded synaptic currents in voltage clamp while blocking inhibitory synapses (**Figure 4A**). The responses were blocked by the AMPA receptor antagonist NBQX (5 μM) and showed paired-pulse facilitation typical of granule cell parallel fiber synapses onto other targets (**Figure 4B**).

**Figure 4.**
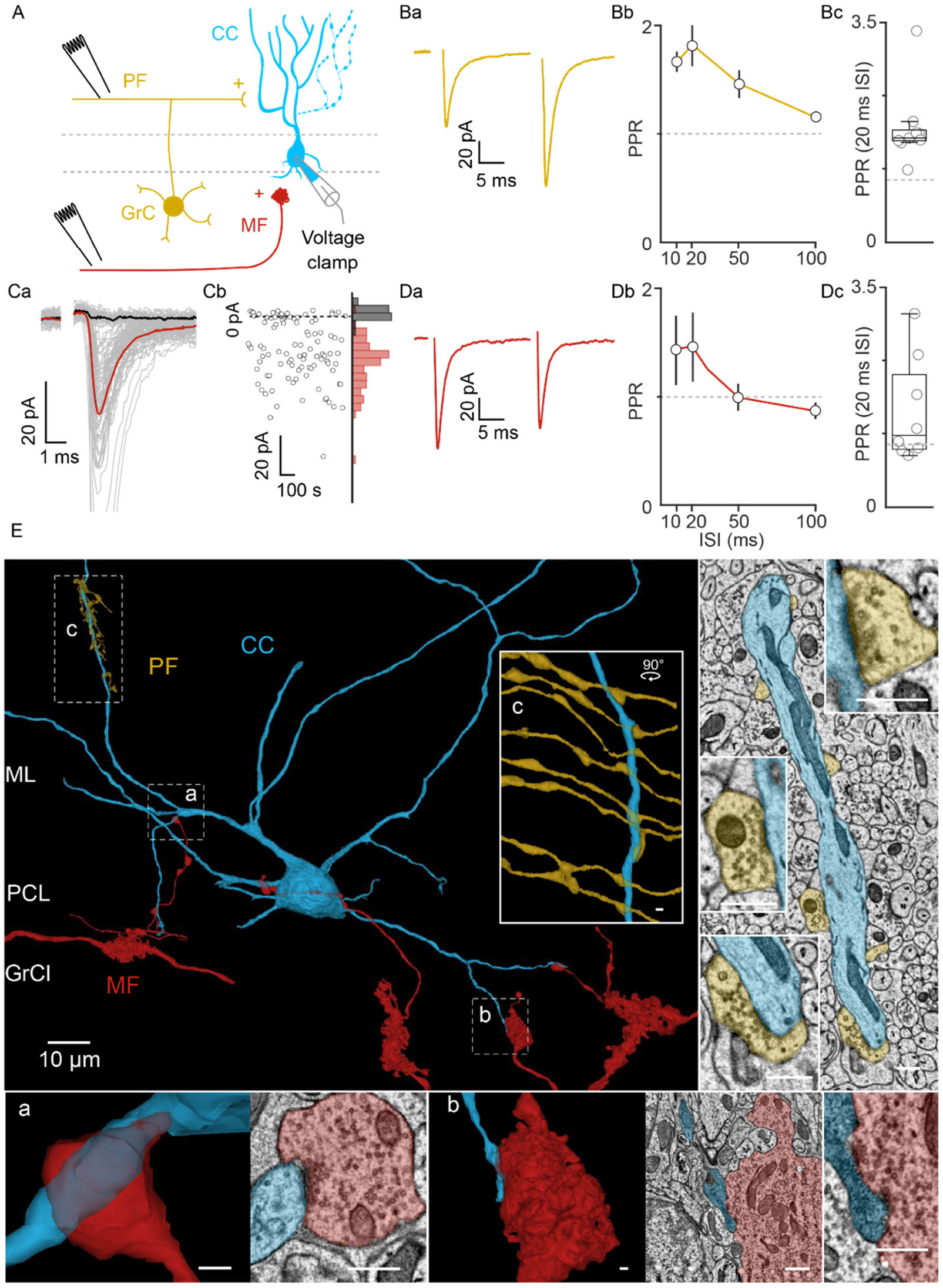
Synaptic excitation of candelabrum cells. A) Schematic of the experimental configuration. EPSCs were recorded from candelabrum cells (CC, blue) in whole-cell voltage clamp during electrical stimulation of GrC parallel fibers (PFs) in the molecular layer, or MFs (red) in the white matter. Ba) Example of facilitating PF-mediated synaptic currents evoked by paired stimulation with an ISI of 20 ms. The Stimulus artifact was subtracted for clarity. Bb) Summary data of paired-pulse ratio of PF-mediated currents evoked at varying ISIs (mean +/- SEM). Bc) Summary of paired-pulse ratio of PF-mediated currents for 20ms ISI. Dashed line indicates PPR=1. Ca) Example of MF-mediated synaptic currents evoked by threshold stimulation. Individual trials (grey), average successes (red) and average failure (black) are shown. The stimulus artifact was blanked for clarity. Cb, left) Amplitudes of synaptic currents in (Ca) as a function of time. Cb, right) Histogram of success and failure trials. Da) Example of non-facilitating MF-EPSC evoked by paired stimulation with an ISI of 20 ms. The stimulus artifact was blanked for clarity. Db) Summary of paired-pulse ratio for MF-EPSCs. Dc) Summary of paired-pulse ratio of MF-EPSCs for a 20 ms ISI. Dashed line indicates PPR=1. E) A CC (blue), and synaptically connected PFs (gold) and MFs (red) reconstructed from serial EM images are shown. Dashed boxes denote regions of insets shown to the right and on the bottom of the figure (a, b, c). a, example of extraglomerular MF synapse onto the basal dendrite of a CC. Left, reconstruction; right, single-plane EM image. b, example of glomerular MF synapse onto the basal dendrite of a CC. Left, reconstruction; right, single-plane EM image. c, PF synapses into the apical dendrite of a CC. Right, single-plane EM images of PF-CC synapses. Inset scalebars: 500nm.

MFs are the primary excitatory inputs into the cerebellar cortex. They form synapses mainly in the granule cell layer where they primarily contact GCs, but also excite GoCs and unipolar brush cells (UBCs), but they do not excite MLIs and PCs whose dendrites are restricted to the molecular layer. Although most CC dendrites are located within the molecular layer, CC cell bodies are located close to the granular layer, and their basal dendrites often extend into the top part of the granular layer (**Figure 3F,3S2**). This raised the possibility that MFs might synapse onto CCs. We tested this by stimulating MF axons in the white matter while blocking inhibitory transmission. White matter stimulation evoked short-latency excitatory currents that were blocked by NBQX (5μM). Stimulation of the white matter at the threshold for MF activation stochastically evoked successes and failures (**Figure 4Ca, Cb**), and slight increases in stimulus intensity eliminated failures. This indicates that this is a single, all-or-none input. We readily identified monosynaptic inputs based on their short latency, because disynaptic MF→GrC→CC inputs had longer, highly-variable latencies. MF inputs also exhibited short-term plasticity that ranged from slight paired-pulse depression to robust facilitation (**Figure 4D**). MF synapses onto granule cells shows a similar variability in paired-pulse plasticity. These results indicate that CCs receive excitatory input from both PF and MFs.

We also assessed connectivity by reconstructing CCs and their inputs in a serial EM volume of adult lobule V cerebellar cortex (Methods, Nguyen et al.2021). We reconstructed all PLIs in the volume and identified CCs based on their characteristic morphological features: soma size, location and shape, the primary dendrite branching in the molecular layer, proximal dendrites extending into the upper granule cell layer, and beaded axons in the molecular layer (**Figure 4E, 4S1**- *light blue*). We also reconstructed PFs and MFs based on their characteristic morphology. (**Figure 4E, 4S1**- *MF:red, PF:gold*). PFs synapsed throughout the entire length of the primary dendrites, and MFs synapsed onto proximal dendrites. MF-CC synapses were of two different types. In some cases a CC dendrite extended into a MF glomerulus and contacted a large MF bouton (**Figure 4E**-*inset b*, **4S1A,B**-*inset b*), but in other cases a small branch from a MF sent a thin axonal projection and contacted CC dendrites (**Figure 4E**-*inset* a, **4S1A**-*inset a*). These reconstructions confirmed that CCs receive synaptic inputs from PFs and MFs.

### Purkinje cells strongly inhibit Candelabrum cells

PCs are spontaneously active at high frequencies, and their collaterals form synapses near the PC layer in the vicinity of CCs (Witter et al., 2016). We hypothesized that PC collaterals could inhibit CCs and provide the high-frequency spontaneous IPSCs we observe in CCs. We used PC-to-CC paired recordings to assess whether such synapses are present. We recorded spontaneous firing of PCs in the loose patch configuration, and measured IPSCs in CCs in voltage clamp (**Figure 5A**). PCs fired spontaneously, and spontaneous IPSCs were observed in CCs (**Figure 5Ba**). Spike-triggered alignment of the traces revealed a short-latency outward current, which was blocked by a GABA_A_R antagonist, consistent with a direct PC-CC connection (**Figure 5Bb, 5C**). PC-CC connections had a moderate success probability (**Figure 5C**, *middle left* and **5Bc**). An individual PC accounted for an average of about 20% of the inhibitory events in a CC (**Figure 5C**, *middle right*), suggesting that a small number of PCs strongly inhibit each CC. This is also consistent with the low probability of finding a directly connected PC-CC pair (8/109 pairs). However, it is likely that some PC collaterals were severed in our experiments, which would reduce the sIPSC frequency and our estimates of the total number of PC inputs.

**Figure 5.**
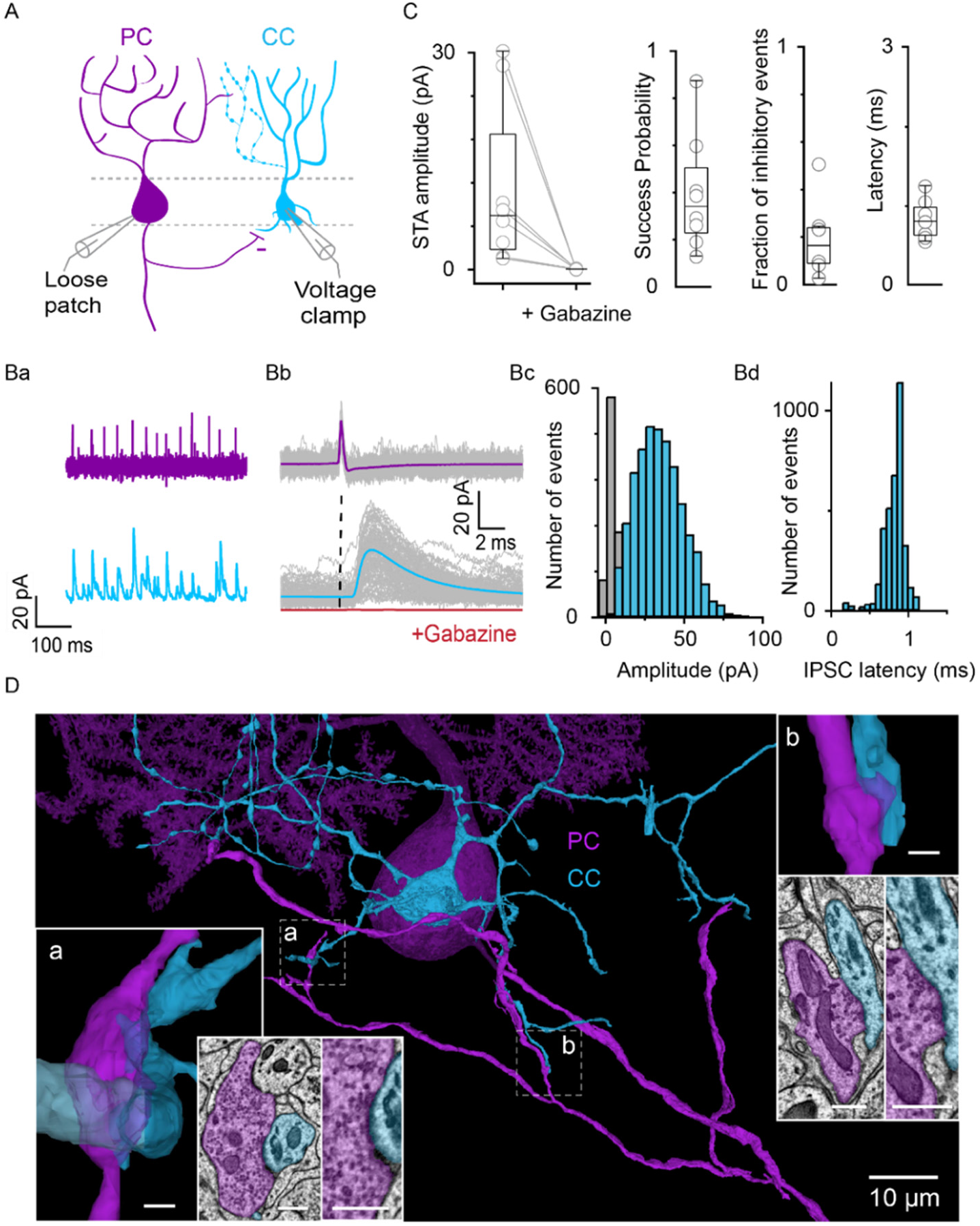
Synaptic Inhibition of Candelabrum cells. A) Schematic of the experimental configuration. IPSCs were recorded from CCs (CC, blue) in whole-cell voltage clamp and spontaneous action potentials were recorded from a synaptically connected PC (purple) in loose-patch configuration. Ba) Loose-patch PC recording (purple) and corresponding whole-cell CC recording in of IPSCs (blue). Bb) Aligned PC spikes (grey: individual spikes, purple: average), and CC recordings (grey: individual traces, blue: average success, dark grey: average failure. Gabazine wash-in eliminated IPSCs in the CC (red trace). Dashed line represents the peak of the action potential. Bc) Amplitude histogram of IPSCs after a PC spike (blue: successes, grey: failures). Bd) Latency histogram of IPSCs relative to PC spike. C) Summary of the properties of PC to CC synapses. n=8/109 pairs. D) EM reconstruction shows a PC (purple) that is synaptically connected to a CC (blue). Dashed boxes denote regions also shown in expanded views along with corresponding EM sections. Scalebars in insets are 500 nm.

We also assessed connectivity using serial EM reconstructions. A PC soma, proximal dendrites and axon collateral are shown (**Figure 5D**, *purple*) along with a CC (**Figure 5D**, light blue). We found that PC axon collaterals directly contact CC dendrites (**Figure 5D, 5S1B**) and somata (**Figure 5S1A**). We found that the same PC forms multiple synapses with a single CC (**Figure 5D, Figure 5S1**). Thus, our EM reconstructions confirmed that PCs directly inhibit CCs.

### Candelabrum cells widely inhibit molecular layer interneurons

The axons of CCs are in the PCL and molecular layer; therefore, their synaptic targets must be cells that have somata or dendrites in the molecular layer. We recorded spiking from CCs in loose patch or current clamp mode, and simultaneously recorded synaptic currents from either MLIs or PCs in voltage clamp (**Figure 6A**). Although we expect CCs to release GABA as a primary transmitter, we did not exclude the possibility of other neurotransmitters; therefore, we omitted synaptic blockers from the bath solution and recorded postsynaptic currents at intermediate voltages between E_CI_- and the reversal potential of EPSCs (0 mV), as shown for a CC-MLI paired recording (**Figure 6B**). Spike-triggered averages showed low latency IPSCs in MLIs (**Figure 6C, E**- *far right*) that were blocked by bath application of GABAzine (**Figure 6C**- *red trace*, **6E**). The amplitude histogram of success and failures is shown for the example pair (**Figure 6D**). We found that 10% (5/50) of CC-MLI pairs were connected, and the properties of these synapses are summarized (**Figure 6E**). Similar experiments were performed to assess CC to PC connections, and we only observed a single connected pair (n= 1/83 pairs, 1%) **(Figure 6F**). It is likely that we underestimate the fraction of cells inhibited by CCs because slicing can sever all or part of the CC axon.

**Figure 6.**
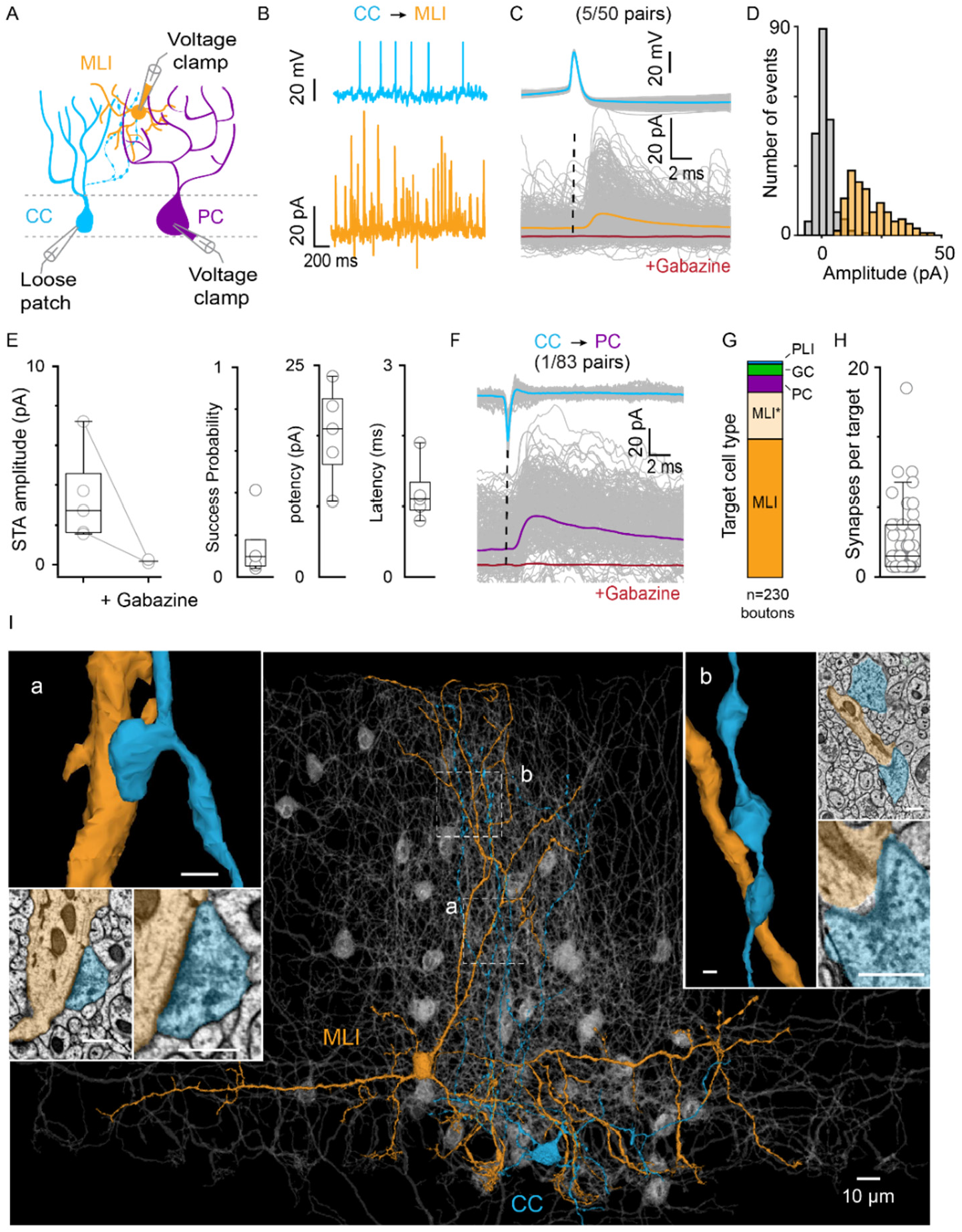
Candelabrum cell inhibition of targets. A. Recording configuration is shown in which spontaneous action potentials were recorded from CCs in loose patch or current clamp, and PCs or MLIs were simultaneously recorded in voltage clamp. B. Raw traces of a CC-MLI paired recording. CC action potentials (top) and voltage clamp recording of MLI (orange). C. Aligned CC action potentials (top, grey: individual traces, blue: average) and MLI traces (bottom, grey: individual traces, orange: average, red: average in the presence of gabazine). Dashed line represents the peak of the action potential. D. Histogram of IPSC amplitudes in C (Orange: successes, grey: failures). E. Summary of properties of CC to MLI connections. n=5/50 pairs. F. Aligned CC action potentials (top, grey: individual traces, blue: average trace) and PC traces (bottom, grey: individual traces, purple: average, red: average in the presence of gabazine). G. Fraction of CC synapses received by different cell types (n=230 boutons). H. Number of CC output synapses received per target cell, from EM reconstructions. I. EM reconstruction of a CC (blue) and its postsynaptic target cells (grey). One target basket cell is highlighted (orange). Dashed boxes denote regions of insets showing example synapses to this cell. a,b CC axon boutons contacting a basket cell dendrite with corresponding single section views of the synapses. Inset scalebars: 500 nm.

We then identified the CC synaptic boutons and their targets in EM reconstructions. We found that most CC synapses are made onto MLIs (197/230= 85.6% of synapses), but CCs also synapse onto PCs (7.8%,18/230), GoCs (5.2%,12/230) and PLIs (1.3%, 3/230) **(Figure 6G).** These findings are in qualitative agreement with our electrophysiological studies that found CCs were more likely to inhibit MLIs than PCs. We found that a single CC can inhibit the majority of nearby MLIs (**Figure 6I**-*MLI grey, CC-light blue*) with a variable number of synapses per target ranging from 1 to >10 (**Figure 6H**). Example high resolution views are shown for a target MLI (**Figure 6I**- *orange, insets a, b*).

The observation that CCs rarely inhibit PCs but inhibit many MLIs, raised the possibility that the primary role of CCs may be to regulate MLI inhibition of PCs. MLIs are spontaneously active and are the primary source of inhibition to PCs. Therefore, anything that alters MLI firing will indirectly influence PCs. We tested the hypothesis by increasing CC firing while simultaneously recording spontaneous inhibition in nearby PCs (**Figure 7A**). These recordings were conducted in the absence of blockers, so the large spontaneous IPSCs are outward, and small excitatory EPSCs are inward. In an example pair, elevating CC firing to 97 Hz for 5s markedly deceased spontaneous inhibition in a nearby PC (**Figure 7Ba**). The firing of single CCs to 60 spikes/s decreased IPSC charge by an average of 24±5% (**Figure 7C**). Thus, a single CC can powerfully disinhibit nearby PCs.

**Figure 7.**
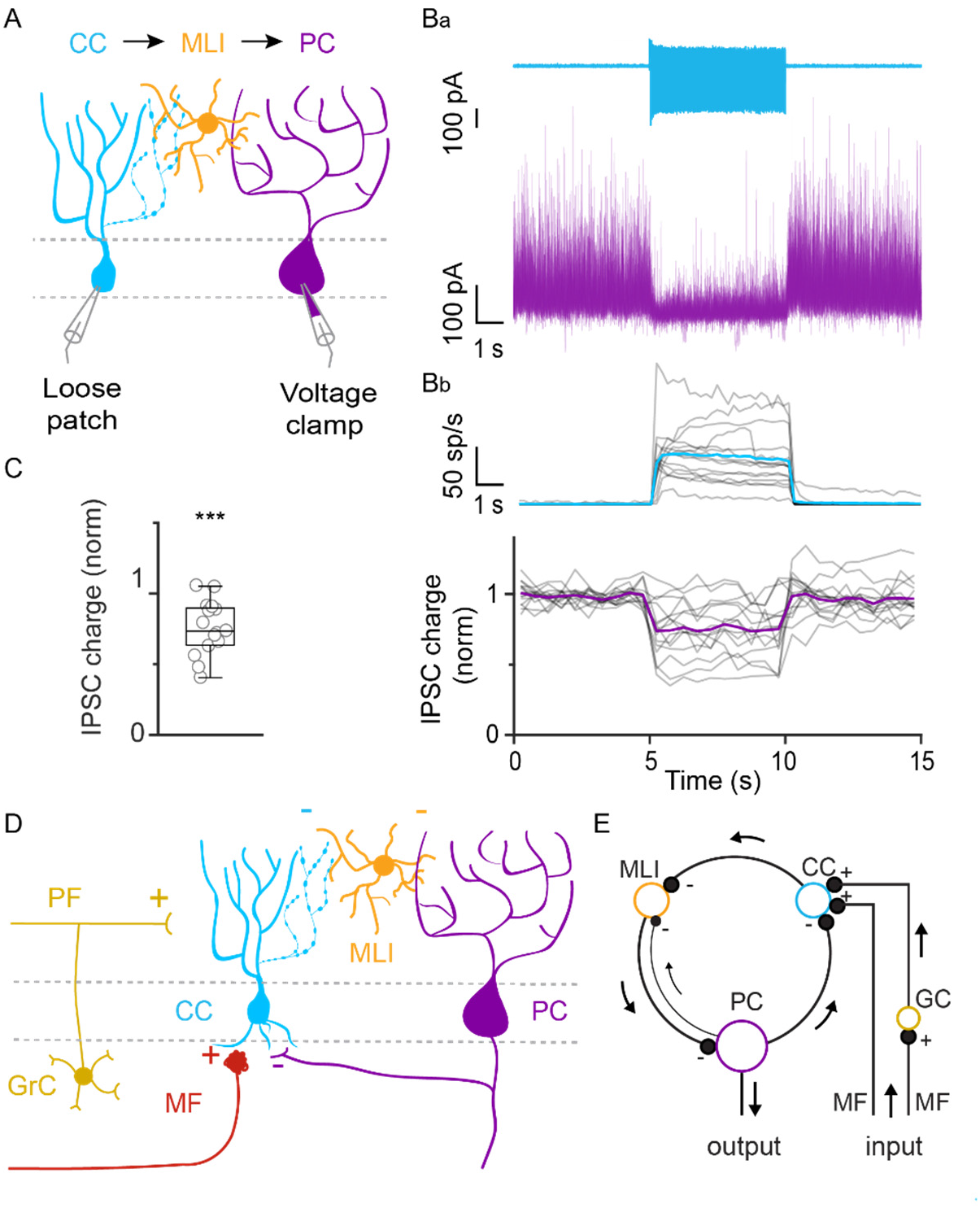
Candelabrum cells disinhibit Purkinje cells. A. Schematic showing experimental configuration of paired recordings in which a CC was stimulated while recording spontaneous IPSCs in PCs. Ba. Elevated CC activity (light blue) decreases spontaneous inhibition of PCs (purple). Bb. Summary of the time course of CC firing and total IPSC charge in PCs. Grey traces show the mean of individual cells (>5 trials) and colored traces show the mean of all cells, p= 3.1e-04, Wilcoxon signed rank test, n=15 pairs. C. Summary the effect of CC excitation on PC inhibition, n=15 pairs. D. Schematic of the circuitry of the CC within the cerebellar cortex. E. Redrawn schematic emphasizing the ring of inhibitory neurons comprised of PCs, CCs, and MLIs and the regulation of CC firing by PC outputs, MF inputs and granule cells.

## DISCUSSION

We show that CCs represent an unexpectedly large population of molecularly distinct interneurons that are present within all regions of the cerebellar cortex. CCs are highly excitable and receive inputs from all processing stages, including MFs, granule cells, and PCs. CCs in turn inhibit MLIs. Because of their unique position within the cerebellar circuit, CCs are predicted to be a critical element for cerebellar processing

### CCs are a distinct population of cells

Whether CCs constitute a distinct cell type was very much an open question. The initial study of CCs described a cerebellar interneuron with a morphology that differed from PCs, GoCs, UBCs and granule cells. Although CCs are similar to MLIs, in that they have dendrites and axons largely restricted to a sagittal plane of the molecular layer, the morphology of CCs differs in important ways. CCs often have dendrites that extend into the granular layer, which is not the case for MLIs. In addition, MLIs with cell bodies near the PC layer tend to make distinctive basket cell synapses onto the soma and axon of PCs, whereas CCs do not. However, the possibility remained that CCs might be an MLI subtype. The relationship of CCs to other PLIs, namely GLCs and LCs, was unknown.

It is remarkable that despite CCs being identified twenty-seven years ago, subsequent studies failed to provide any insight into the properties and connectivity of CCs within the cerebellar circuit. Several factors had hindered the study of CCs. First, even though they are abundant relative to PCs and GoCs, their numbers are dwarfed by the much more numerous granule cells that are located nearby. Second, even if recordings were restricted to fluorescently labelled inhibitory neurons, CCs would be obscured by nearby PCs that are much larger than CCs. Third, the application of single cell molecular approaches to the study of the CCs had also been limited by granule cells outnumbering most other cell types, especially interneurons (Saunders et al., 2018). Consequently, initial studies of the cerebellar cortex were unable to provide much insight into CCs and other PLIs. We overcame the challenge of profiling cells in the cerebellar cortex by transcriptionally profiling 780,553 nuclei, which allowed us to identify three populations of PLIs. Nonetheless it was not possible to link the molecular properties of PLIs to CCs until we found that neurons that matched the original description of CC morphology were labelled in Oxtr-cre x ai14 mice (**Figure 3F, 3S2)**. This allowed us to fill cells to determine their morphology, determine their electrophysiological properties, study their synaptic connections, and link these properties to the molecular identity of these cells.

Our pseudotime logistic regression analysis of differential gene expression showed that CCs are molecularly distinct from MLIs, GoCs and LCs (**Figure 1D**), but that there are some similarities in the molecular properties of CCs and GLCs. The degree of molecular similarity of CCs and GLCs exist is similar to that MLI1 subtypes. Nonetheless there is a rather sharp distinction in the expression of *Slc6a5*, suggesting that GLCs release both glycine and GABA, but CCs exclusively release GABA. Oxtr-Cre mice allowed us to target CCs, rather than glycinergic GLCs. There are also similarities and differences in the locations, morphologies and connections of CCs and GLCs. The cell types are both located near the PC layer, although the locations of CC somata extend slightly into the molecular layer, and GLCs extend into the granular layer. GLCs and CCs are both inhibited by PCs (**Figure 5**). However, CCs and GLCs have distinct dendritic and axonal morphologies. CC axons ascending locally in a sagittal plan whereas GLC axons extending in a transverse orientation (Hirono et al., 2012; Laine and Axelrad, 2002; Schilling et al., 2008). Additionally, CCs differ in electrophysiological properties in that they have much higher input resistance and are more excitable than GLCs (Hirono et al., 2012). Thus, differences in gene expression, morphology and electrophysiological properties indicate that CCs constitute a distinct type of cerebellar interneuron.

It is important to recognize that even with the addition of the CC as a distinctive type of inhibitory interneuron, the circuitry of the cerebellar cortex is comprised of a small number of cell types. There are just seven types of inhibitory interneurons in the cerebellar cortex: MLI1 (MLI1_1 and MLI1_2 subtypes), MLI2, Golgi1, Golgi2, CC, LC, and GLC. This contrasts with other brain regions, such as the cerebral cortex, where there are 42-59 distinct GABAergic types in M1 (Adkins et al., 2020; Bakken et al., 2020; Yao et al., 2020), or the CA1 region of the hippocampus, where more than 20 distinct types of GABAergic interneurons have been identified (Pelkey et al., 2017; Qian et al., 2020). This suggests that the characterization of a new type of interneuron in the cerebellum is of particular significance.

### CCs are widespread and prevalent

CCs were found in all regions of the cerebellar cortex. Our smFISH results show that CCs are present in all lobules of the vermis of male and female mice. In addition, our initial transcriptional profiling was based on the analysis of 16 regions of the cerebellar cortex that included regions in the vermis, lateral lobules, the flocculus and the paraflocculus, and CCs were present in all regions (**Figure 3 S3**).

CCs are also a prevalent type of interneuron. We found that there were approximately 80% as many CCs as there were Golgi1 cells (based on smFISH). Moreover, CCs are the most abundant PLI with approximately 43% more CCs than GLCs (based on smFISH).

### Candelabrum cells within the cerebellar circuit

Our studies indicate that PCs are the primary source of inhibition to CCs. Paired recordings established that a single PC can account for up to 50% of the total inhibitory events in a CC (mean=19%), and that the PC-to-CC input has a moderate success probability of approximately 0.4. We never observed inhibition from MLIs to CCs in paired recordings (n=0/15 pairs), but we found PCs typically made multiple synapses onto a CC. Together, these findings suggest that a modest number of PCs powerfully inhibit each CC.

CCs receive extensive excitatory inputs from many granule cells, as expected for a neuron with dendrites in the molecular layer. We also found that MFs provide rapid short latency excitation. This was somewhat surprising, because MFs are generally thought to be restricted to the granular layer, and CC dendrites only extend slightly into the granular layer. Serial EM reconstruction revealed that CCs receive input from multiple MFs (**Figure 4, 4S1**) in the form of glomerular and extraglomerular synapses.

Our electrophysiological and molecular studies established that CCs are GABAergic cells. CC synapses onto MLIs and PCs were examined under conditions that allowed us to assess the contribution of other transmitters including glutamate, GABA and glycine. Synaptic currents were eliminated with a GABA_A_ receptor antagonist, confirming that they are exclusively GABAergic and lack a glycine component. This is consistent with the lack of *Slc6a5* that encodes the glycine transporter Glyt2 that is expressed by glycinergic neurons such as GLCs and LCs.

### Possible functional relevance and significance

CCs are part of a primarily unidirectional inhibitory loop comprised of PCs, CCs and MLIs that has intriguing properties (**Figure 7D,E**). This simple circuit has the property that whenever the firing rate of any element in the loop is altered, this circuit will counteract that change in firing rate. In addition, such ring oscillators with an odd number of elements have been studied extensively as a means of generating oscillations that in some cases decay exponentially in amplitude (Friesen and Stent, 1977; Horikawa, 2011). The nature and duration of these oscillations depend on many factors. Functionally, ring oscillators have the potential to allow a circuit of exclusively inhibitory neurons to generate oscillations, and such a circuit also has the potential to generate undesirable epileptiform activity. Transient cerebellar oscillations sometimes precede specific actions, and can be coincident with oscillations in other brain areas, like motor cortex. It has been proposed that that gap-junction coupled Golgi cells could enable such oscillations (Vervaeke et al., 2010). The circuit properties we describe here for CCs suggest that they could also generate functionally-relevant oscillations following MFs and GrC activation and engagement of the CC to MLI to PC loop.

The synaptic inputs and outputs set CCs apart from the better characterized cerebellar interneurons, GoCs and the MLIs (**Figure 7D**). CCs are excited by MFs, powerfully inhibited by PCs, and they directly inhibit cells in the molecular layer. Although GoCs are also excited by granule cells and MFs, they do synapse onto cells in the molecular layer and are not inhibited by PCs. MLIs inhibit cells in the molecular layer and some are inhibited by PCs, but MLIs are not excited by MFs. Thus, only CCs simultaneously sample multiple stages of cerebellar processing, including the MF inputs, PC outputs and granule cells. CCs are therefore important circuit elements positioned within the circuit if the cerebellar cortex to weigh the activity of inputs, outputs and within the cerebellum to control the output of the cerebellar cortex.

It is more difficult to compare the circuit properties of CCs to other PLIs, because the synaptic connectivity of LCs and GLCs has not been comprehensively described. Both of these cell types reside near the PCL where PC collateral synapses are abundant, so it is not surprising that they are also strongly inhibited by PCs. Other PLIs may also be similar to CCs with regard to receiving MF inputs. Stimulation of the granular layer evoked depressing responses in GLCs, which may reflect MF inputs (Hirono et al., 2012). Light level immunofluorescence also suggested that LCs receive MF inputs (Miyazaki et al., 2020) and suggest that they may also inhibit nearby MLIs. However, LCs differ from CCs in that they powerfully inhibit GoCs with glycinergic/GABAergic synapses, whereas CC to GoC synapses are exceedingly rare. In addition, LCs and GLCs have much longer axons than CCs, that allow them to influence targets at much further distances than CCs. Thus, it seems as if CCs and other PLIs have some common circuit properties and some unique specializations, but a more complete comparison is not possible until more is learned about the connectivity of LCs and GLCs.

CCs can exert a large influence on PC excitability, and can be thought of as master regulators of PC excitability that are privy to input, output and intrinsic activity. We have shown that a single CC can dramatically alter the inhibition that PCs receive from MLIs, with up to a 60% reduction of inhibitory charge (**Figure 7**). EM reconstructions confirmed the physiology experiments, but also revealed that CCs inhibit the majority of nearby MLIs. GCs and MFs promote CC firing, reduce MLI firing and disinhibit PCs. This is countered by the activity of PCs themselves, which will have the opposite effect. When PC firing is elevated, it will decrease the firing of CCs that in turn increases the firing of MLIs, thereby resulting in negative feedback to the PC that counteracts the increase in CC firing. These properties, along with the observation that CCs are abundant and widespread, indicate that the properties of CCs must be considered in order to fully appreciate the manner in which the cerebellar cortex functions and contributes to behaviors.

## CONTACT FOR REAGENT AND RESOURCE SHARING

Further information and requests for resources and reagents should be directed to and will be fulfilled by the Lead Contact, Wade Regehr.

## EXPERIMENTAL MODEL AND SUBJECT DETAILS

### Animals

Animal procedures have been carried out in accordance with the NIH and Animal Care and Use committee (IACUC) guidelines, and protocols approved by the Harvard Medical School and Broad Institute Standing Committee on Animals.

## METHOD DETAILS

### Electrophysiology

#### Slice preparation

Juvenile animals of either sex aged 28-45 days old were anesthetized with an intraperitoneal injection of Ketamine (10 mg/kg) and then perfused transcardially with ice cold cutting solution containing (in mM) 110 CholineCl, 7 MgCl_2_, 2.5 KCl, 1.25 NaH_2_PO_4_, 0.5 CaCl_2_, 25 Glucose, 11.5 Na-ascorbate, 3 Na-pyruvate, 25 NaHCO_3_, equilibrated with 95% O_2_ and 5% CO_2_. The brain was rapidly dissected, and the cerebellum was cut into 250 μm thick parasagittal slices in the same solution on a vibratome (VT1200S, Leica). Slices were then transferred to 34°C warm artificial cerebrospinal fluid (ACSF) containing (in mM) 125 NaCl, 26 NaHCO_3_, 1.25 NaH_2_PO_4_, 2.5 KCl, 1 MgCl_2_, 1.5 CaCl_2_, and 25 glucose, equilibrated with 95% O_2_ and 5% CO_2_ and incubated for 30 min. Slices were then stored at room temperature until recording for up to 6 hours.

#### Recordings

CC, MLI and PC recordings were performed at ~32 °C with an internal solution containing (in mM) 150 K-gluconate, 5 KCl, 10 HEPES, 10 MgATP, 0.5 GTP, 5 phosphocreatine-tris2, and 5 phosphocreatine-Na2, and 0.1 Alexa 594 (pH adjusted to 7.2 with KOH, osmolality adjusted to 310 mOsm/kg). The chloride reversal potential was adjusted to −85 mV. Visually guided whole-cell recordings were obtained with patch pipettes of ~4 MΩ resistance pulled from borosilicate capillary glass (BF150-86-10, Sutter Instrument, Novato, CA). Electrophysiology data was acquired using a Multiclamp 700B amplifier (Axon Instruments), digitized at 20 kHz and filtered at 4 kHz. For isolating inhibitory synaptic currents in voltage clamp and the following receptor antagonists were added to the solution (in μM): 2 R-CPP, 5 NBQX, 1 strychnine, 1.5 CGP. All drugs were purchased from Abcam (Cambridge, MA) and Tocris (Bristol, UK). To obtain an input-output curve, CCs were injected with a constant hyperpolarizing current to hold them at ~−65 mV, and 500 ms long current steps ranging from −30 pA to +30 pA were injected in 5 pA increments. Capacitance and input resistance (Ri) were determined using a 10 pA, 50 ms hyperpolarizing current step. Spontaneous firing rate and resting membrane potential were determined using cell-attached recordings in voltage clamp mode. To determine Vm, a series of voltage ramps from +100 to −120 mV were averaged, and the reversal potential of the leak current was calculated by the interception of a linear fit to the leak current and the voltage activated potassium current (Verheugen et al., 1999). For the experiments in Figure 7, CC firing was evoked in loose-patch mode in voltage clamp by stepping the pipette voltage to positive values (range +70 to +150mV).

For measurement of the chloride reversal potential, we recorded CCs in the presence of NBQX, CGP, CPP and strychnine using gramicidin perforated patch. We recorded spontaneous IPSCs at voltages between −95mV and-60mV, measured the sIPSC charge for each voltage and calculated the reversal of the sIPSC charge. Traces were low-pass filtered with a Savitzky-Golay filter (order 3, frame= 501) and slow fluctuations were removed by subtracting a median filtered version of each trace (kernel=10000=0.5s) prior to sIPSC charge measurement. The internal solution was 110 K-methanesulfonate, 13 NaCl, 2 MgCl_2_, 10 EGTA, 1 CaCl_2_, 10 HEPES, 2mM QX314 and 10μg/ml of Gramicidin A (Sigma-Aldrich).

Acquisition and analysis of electrophysiological data were performed using custom routines written in MATLAB (Mathworks, Natick, MA) and IgorPro (Wavemetrics, Lake Oswego, OR). Data are reported as median ± interquartile range for box plots and as mean for bar graphs, and statistical analysis was carried out using the Wilcoxon signed rank test or a Poisson generalized linear mixed effect model of the form SpikeCount ~ 1 + CurrentSteps + CellType + CurrentSteps*CellType with a canonical log link function, fitted using maximum pseudo-likelihood estimation, as indicated. Statistical significance was assumed at p < 0.05.

### 2-photon imaging and analysis

CCs were filled with 100μM Alexa-594 through the patch pipette to visualize their morphology using 2-photon imaging. We used a custom-built 2-photon laser scanning microscope with a 40X water immersion objective (Olympus, 0.8 NA) and a pulsed 2-photon laser tuned to 800 nm (Mira 900, Coherent, Santa Clara, CA). We acquired Z-stacks with 1 μm Z-step of Alexa-594 fluorescence and LDIC. 2 photon images were processed in ImageJ to create maximum intensity Z projections of Alexa-594 fluorescence which were subtracted from a single DIC focal plane.

### Floating Slice Hybridization Chain Reaction on fixed cerebellar slices

#### Tissue preparation

Male/female p44-45 mice were anesthetized with an intraperitoneal injection of ketamine (10 mg/kg) and transcardially perfused with ice-cold PBS, followed by ice-cold 4% PFA in PBS. Brains were postfixed overnight in 4% PFA at 4 °C, washed in PBS, then sliced sagittally at a thickness of 40μm using a vibratome (VT1000S, Leica) and transferred to a solution of 70% ethanol in RNAase-Free water until HCR was performed.

#### HCR

A floating slice HCR protocol was performed (based on <u>dx.doi.org/10.17504/protocols.io.bck7iuzn</u>) with the following HCR probes and hairpins from Molecular Instruments, Inc. (Los Angeles, CA): Neurexophilin 1 (*Nxph1*), Aldehyde dehydrogenase 1 family, member A3 (*Aldh1a3*) and Sodium-and chloride – dependent glycine transporter 2 (*Slc6a5*). Amplification hairpins were B1-647, B2-488 and B3-546.

#### Imaging

Slices were mounted on a glass slide with ProLong Glass mounting medium without DAPI (ThermoFisher Scientific) and imaged on a widefield slidescanner (Olympus VS120) using a 40X air objective (Olympus, 0.95 NA) and a 16-bit camera (Hamamatsu Orca-R^2^ model C10600) with 7-11 focal planes (3μm per Z plane) per channel and automatic tiling. Focusing was done manually for ~100 focal reference points throughout each slice.

#### Image processing

Image processing was done in Matlab R2019b (Mathworks), unless otherwise specified. Background fluorescence was reduced by subtracting a median filtered version for each individual focal plane (kernel size= 20 x 20 px). After this, puncta were detected using Fast 2D peak finder (Matlab central file exchange, Natan 2020) with a threshold determined by the median absolute deviation of the image, then masked with a circular kernel (radius= 10px) and Z projected as a maximum intensity projection. Each channel was processed separately.

Max Z projections were downscaled to half resolution and registered to a reference slice using a geometric transformation fit to control point pairs (4^th^ order polynomial transformation). Control points were manually selected along the PCL and edge of the molecular layer.

Subsequently, a mask was created for each registered channel by applying a maximum filter (circular kernel, radius= 5 px), followed by a 2D Gaussian filter (sigma=8 px) and manual thresholding. Mask combinations were used to create cell type masks (e.g., *Nxph1*-mask ^ *Aldh1a3*-mask ^ *Slc6a5*-mask = GLC mask), which were used on the background subtracted Z projections. Images masked for each cell type mask were used to manually label the location cells in the slice in ImageJ.

Background fluorescence for the *Nxph1* channel was used to create polygons that delineate the edge of the molecular layer and the PC layer, which were used to measure the distance of each cell to the PCL and Pia. For computing the density of cells in each lobule (**Figure 3D**), cell counts were normalized to the length of the PCL in each lobule. For the layer distribution histograms (**Figure 3E**), cell counts were normalized to the number of slices counted.

In experiments with Oxtr-Cre x Ai14 mice, crosstalk signal originating from tdTomato fluorescence was corrected by subtracting a scaled version of the tdTomato channel from the *Slc6a5* and *Aldh1a3* channels prior to puncta detection. The scaling factor was determined by the average ratio of tdTomato/*Slc6a5* or tdTomato/*Aldh1a3* pixel values from regions in the molecular layer that expressed tdTomato but were devoid of fluorescent puncta.

### snRNA-seq

snRNA-seq data for cerebellar interneurons was obtained in Kozareva et al.2020. Briefly, high throughput single-nucleus RNA-seq (snRNA-seq) was used to analyze 780,553 nuclei isolated from 16 different lobules from 6 p60 mice. Of these, 1177 were categorized as CCs, 735 as GLCs, 531 as LCs, 32716 as MLI1, 10608 as MLI2, 3075 as Golgi1 and 914 as Golgi2.

### Serial Electron microscopy

We previously described the procedures of mouse tissue preparation (Cheadle et al., 2020), sectioning, collection, and TEM imaging (without post-sectioning staining) (Cheadle et al., 2020; Phelps et al., 2021). We used an segmentation workflow based on U-Net CNN to automatically generate the neuron boundary and reconstruct the 3D neuron morphologies (Funke et al., 2019). To deploy the algorithm across hundreds of workers concurrently, we used the Daisy scheduler (https://github.com/funkelab/daisy). To rapidly reconstruct and correct errors from automated segmentation, we developed and used a proofreading work-flow called Dahlia where segmentation is divided into independent cubes; annotators would then select a fragment of a neuron and grow it into the adjacent block until completion. During each growth step, the neuron was visualized and checked for possible split or merge errors.

To identify CCs, we first screened all the small neurons within +-20μm of the center of the PCL, discarded all the GrCs and MLIs, based on their characteristic morphology and focused on PLIs that matched closely the description of CCs, as well as our 2-photon fills. We restricted our analysis to the most completely reconstructed cells, prioritizing the presence of a beaded axon in the molecular layer. Synapses and cell labels were manually generated.

## DATA AND SOFTWARE AVAILABILITY

### AUTHOR CONTRIBUTIONS

TO, SR and W.G.R designed experiments. TO and SR performed electrophysiology experiments. T.N. generated the automated segmentations in the serial EM dataset. N.N. performed smFISH experiments. TO analyzed electrophysiology, smFISH and serial EM data, V.K. and E.M. analyzed snRNAseq data. TO, SR and W.G.R wrote the paper with inputs from all authors.

## ACKNOWLEDGMENTS

We thank members of the Regehr lab and Gord Fishell for comments on the manuscript. This work was supported by grants from the NIH, R01NS032405 and R35NS097284 to W.G.R, NIH/NIMH Brain Grant 1U19MH114821 to E.Z.M., the Stanley Center for Psychiatric Research and by the Vision Core and NINDS P30 Core Center (NS072030) to the Neurobiology Imaging Center at Harvard Medical School.

## Figures and Captions

**Figure 1 S1.**
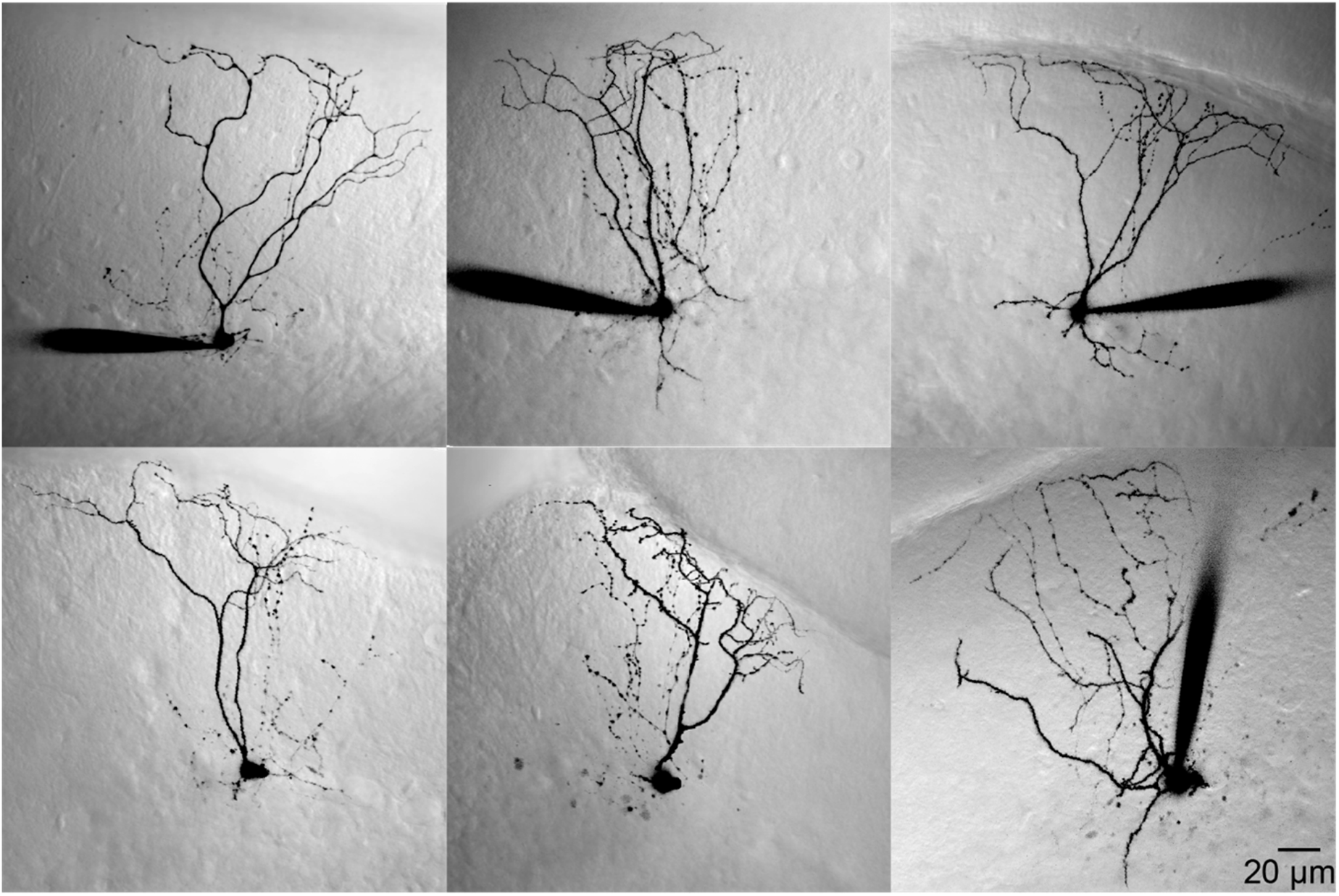
Examples of candelabrum cell morphology. tdTomato-positive candelabrum cells filled with an Alexa594-containing patch pipette and imaged using 2-photon microscopy. Images show LS-DIC in gray indicating the cell location in the slice, and Alexa-594 fluorescence in black as a maximum intensity Z-projection. The pipette was withdrawn prior to imaging for two of the cells.

**Figure 3 S1.**
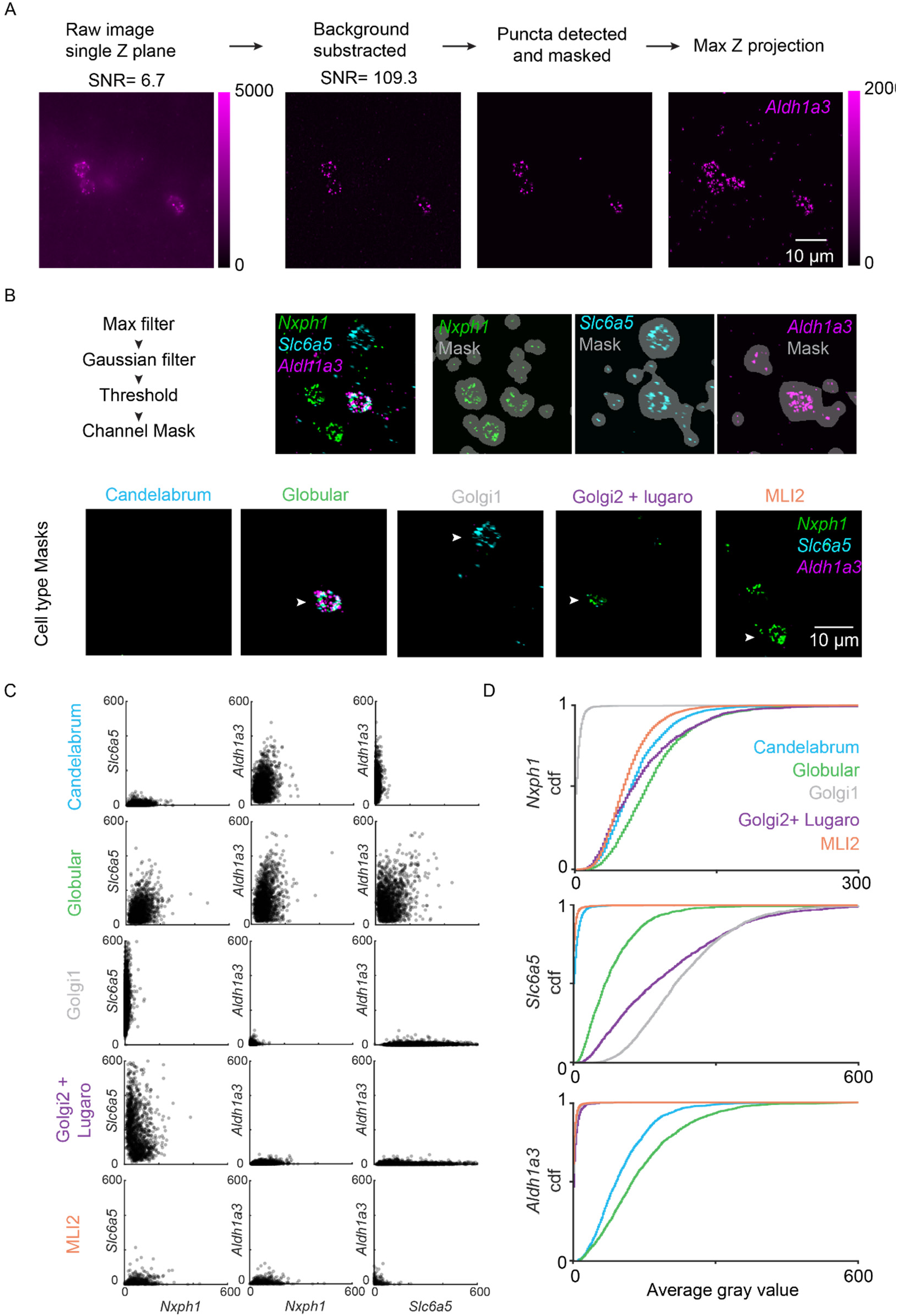
Methods used to analyze FISH data. a. Raw fluorescence image for one channel → image with background fluorescence subtracted→ individual puncta are then detected in each slice→ the results from all slices in a Z-stack are combined. b. Top) a mask is created for each channel. Bottom) Channels masks are combined to create cell type masks. Examples for the same field of view are shown for all cell types. c. Scatter plots of FISH fluorescence are shown for all cells of each cell type. d. Cumulative histograms are shown for *Nxph1, Slc6a5, Aldh1a3* for the different classes of cells.

**Figure 3 S2.**
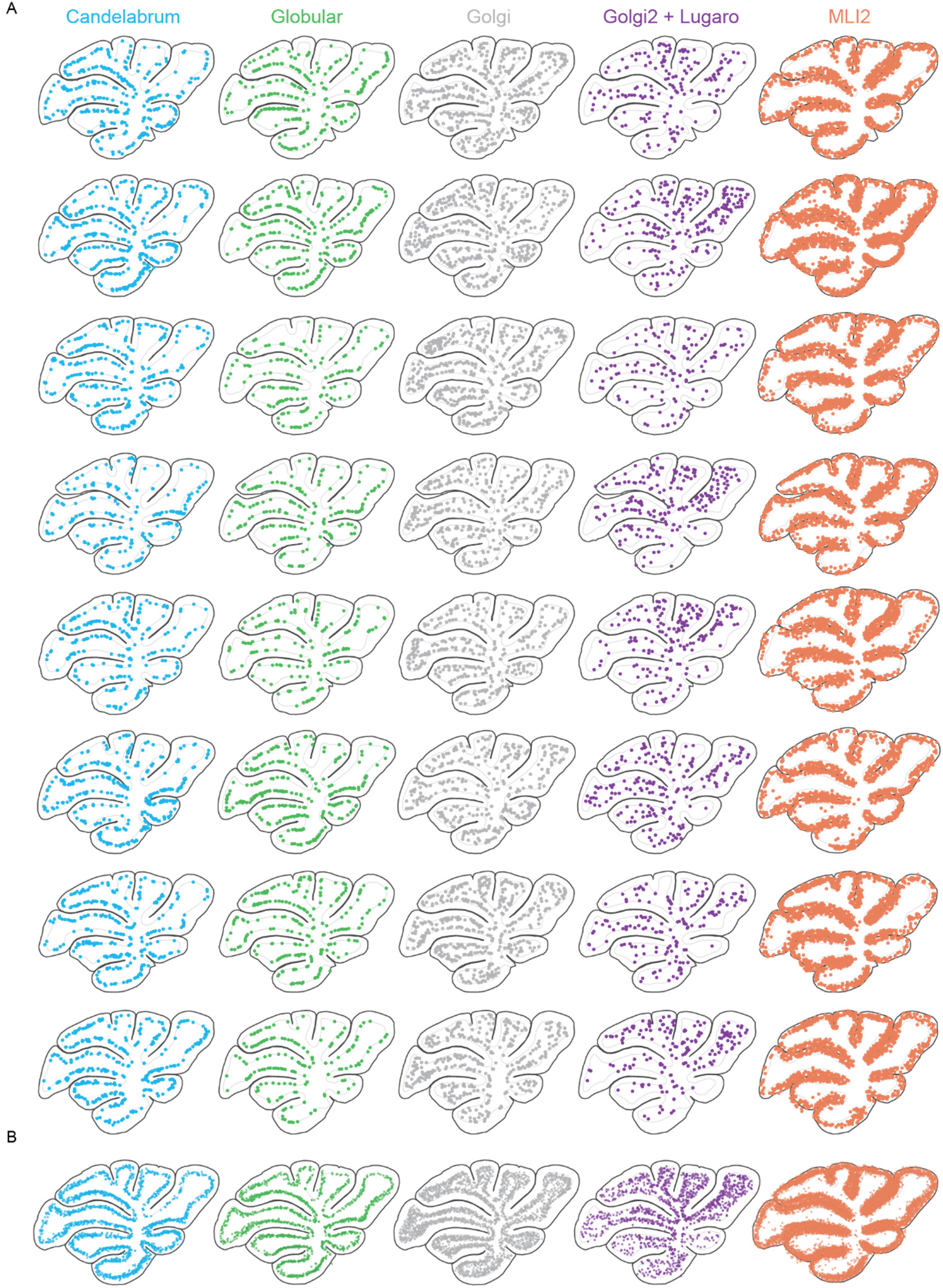
Summaries of locations of five classes of cells in different slices, as in Figure 2. Candelabrum: light blue Nxph1+ Slc6a5- Aldh1a3+ Globular: green Nxph1+ Slc6a5+ Aldh1a3+ Golgi1: grey Nxph1- Slc6a5+ Aldh1a3- Golgi2+ Lugaro: purple Nxph1+ Slc6a5+ Aldh1a3- MLI2: orange Nxph1+ Slc6a5- Aldh1a3- a. Four slices each from a female (top) and a male (bottom) are shown. b. The locations of cell types from all slices are overlayed. The symbol sizes have been reduced to allow better visualization of the locations of individual cells.

**Figure 3 S3.**
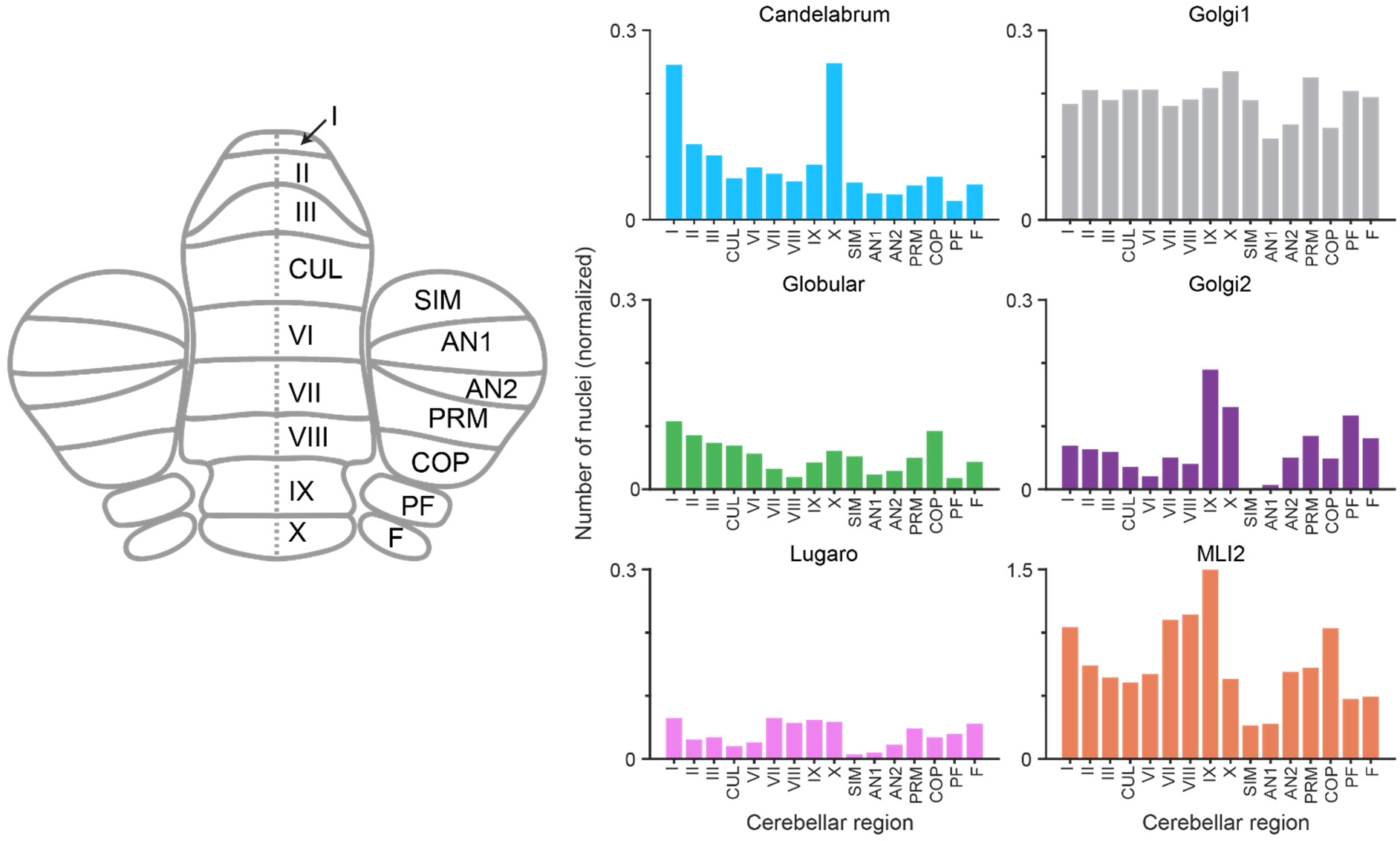
snRNAseq nuclei counts for Candelabrum, Globular, Lugaro, Golgi1, Golgi2 and MLI2 cells from different regions of the cerebellum. Nuclei counts are normalized to the number of PC nuclei in each region. CUL=Culmen, SIM=lobule simplex, AN1= Crus I of ansiform lobule, AN2= Crus II of ansiform lobule, PRM= paramedian lobule, PF=paraflocculus, F=flocculus. Data are from Kozareva et al. 2020.

**Figure 3 S4.**
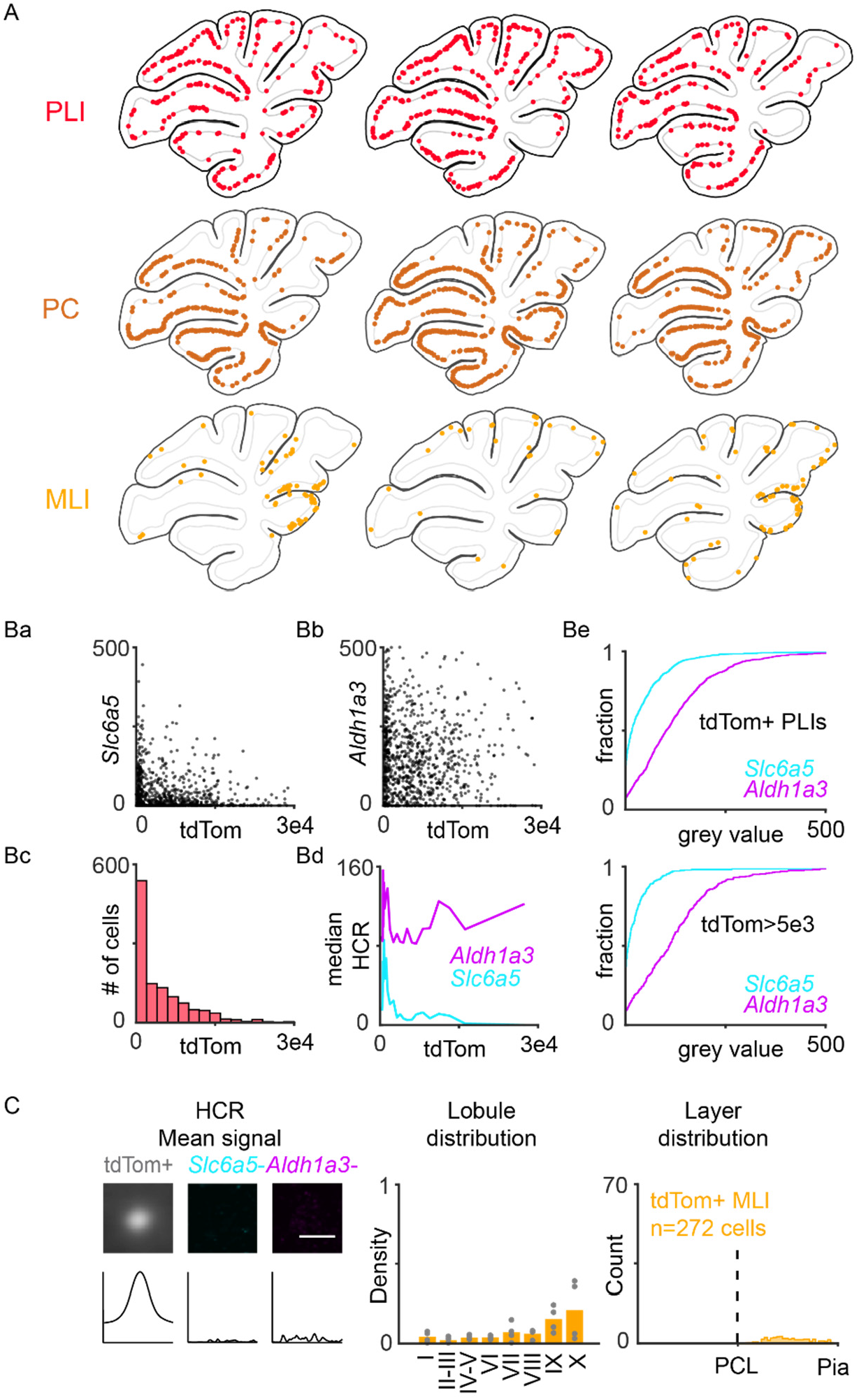
Properties of labelled cells in Oxtr-Cre x Ai14 mice. A. Slice location of tdTomato labeled cell types for different cerebellar slices. Small PCL cells (top, n=1206 cells), PCs (middle, n=1862 cells) and some MLIs (bottom, n=272 cells) were labeled. Ba) Scatter plot of *Slc6a5* signal for individual PLIs as a function of TdTomato fluorescence. Bb) Scatter plot of *Aldh1a3* signal for individual PLIs as a function of TdTomato fluorescence. Bc) Histogram showing distribution of tdTomato fluorescence intensities in PLIs. Bd) Median HCR signal for PLIs as a function of TdTomato fluorescence. Be) Cumulative plots showing distribution of HCR mean values in all tdTomato expressing PLIs (top) and in PLIs with high tdTomato fluorescence (bottom). C. Summary showing sparse tdTomato labelling of MLIs in Oxtr-Cre x ai14 mice. MLIs do not express *Slc6a5* or *Aldh1a3*. These cells exist in low density in the most frequent recording regions (lobule VI-VIII). Lobule IX and X were avoided for electrophysiology recordings because they showed the highest density of MLI labeling.

**Figure 4 S1.**
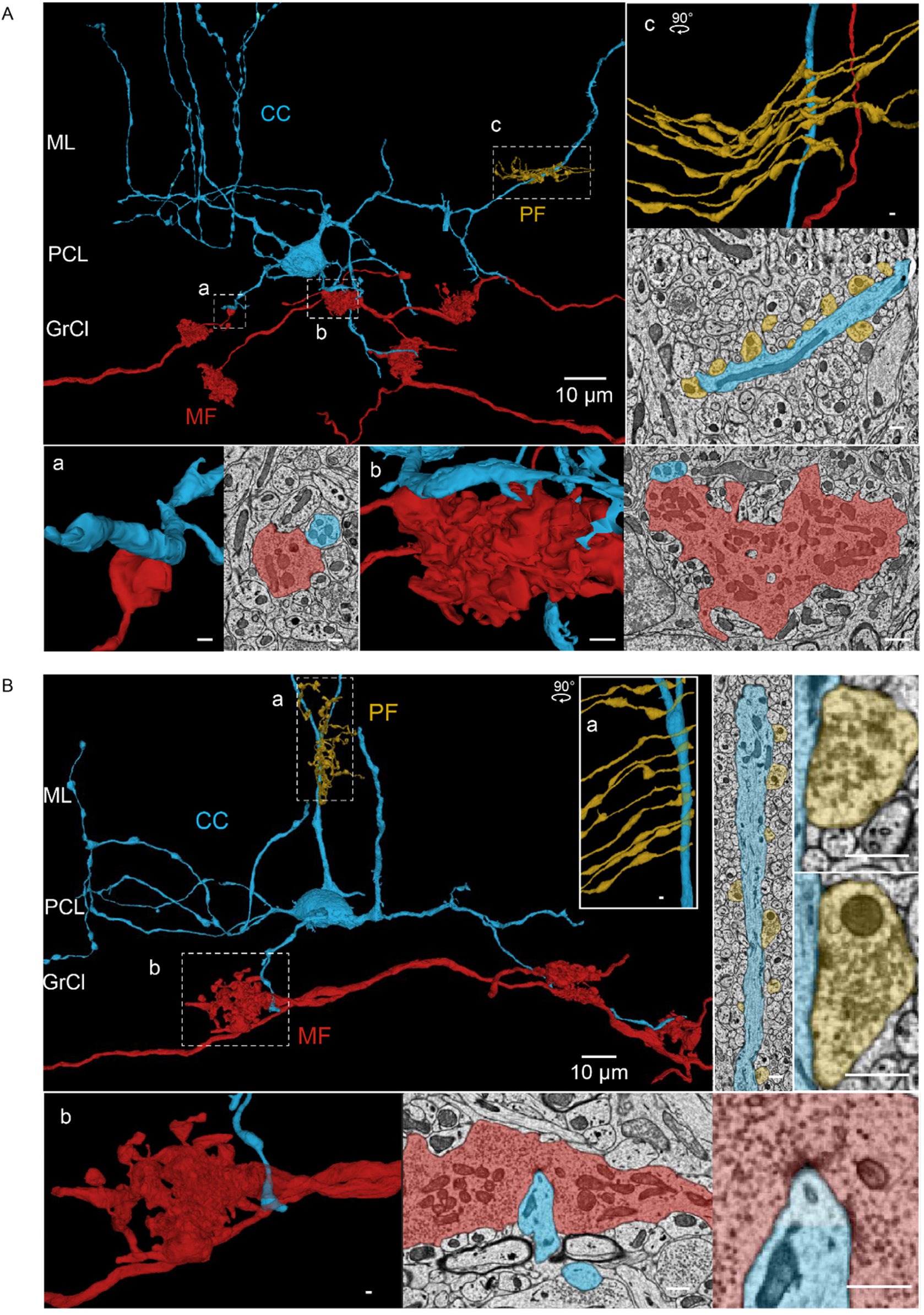
Examples of MF (red) and granule cell PF (yellow) excitatory inputs to CCs (blue) for two different cells (A and B). inset scalebars: 500 nm.

**Figures 5 S1.**
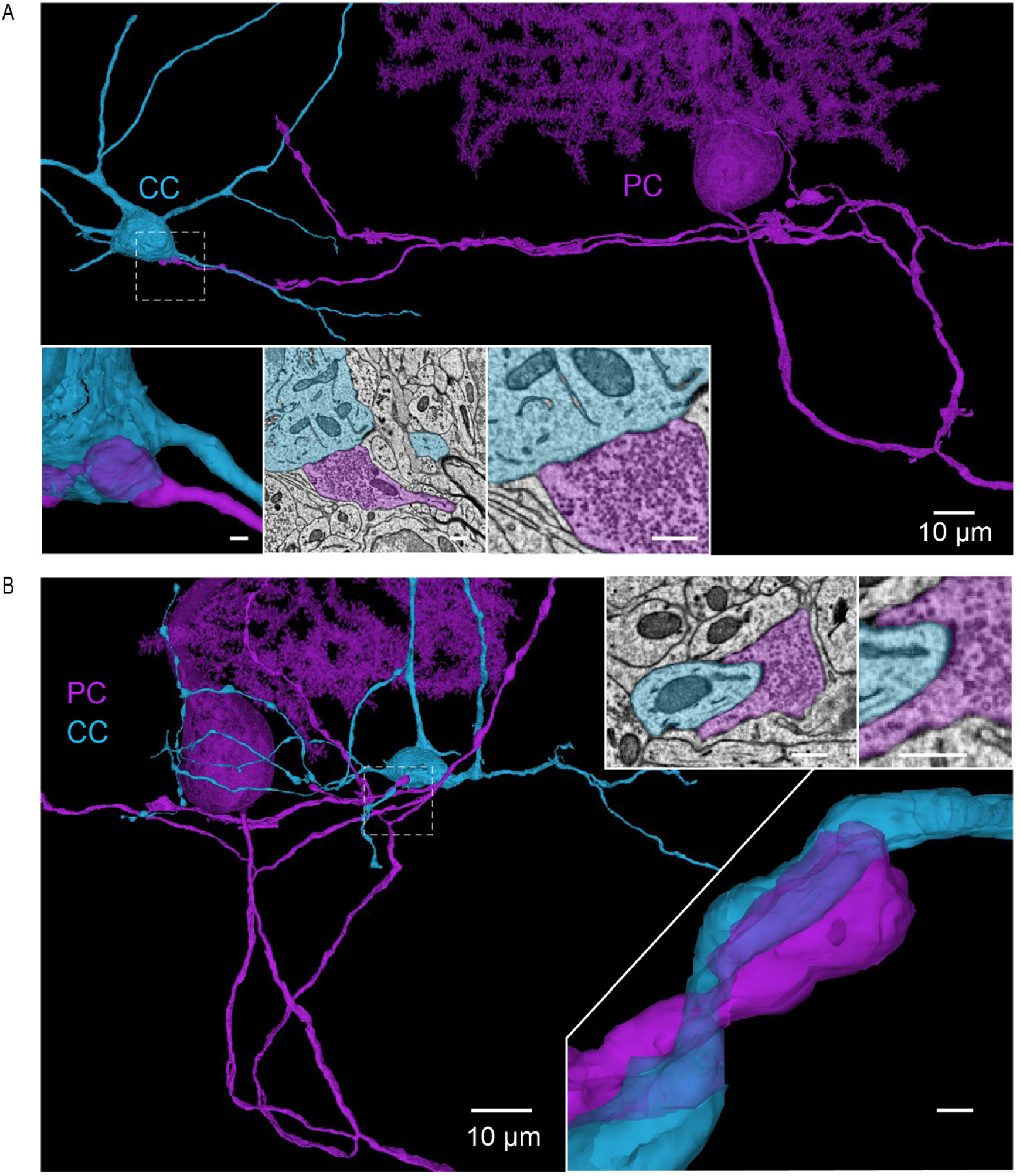
Additional examples of PC (purple) inhibition of candelabrum cells (light blue). Inset scalebars: 500 nm

